# The gut microbiome facilitates ecological adaptation in an invasive vertebrate

**DOI:** 10.1101/2020.12.10.418954

**Authors:** Carla Wagener, Nitya Prakash Mohanty, John Measey

**Author notes:** **Corresponding author:** Carla Wagener. **Ethics:** Collections in Cape Town occurred as part of an ongoing eradication programme (Davies et al., 2020a; Davies et al., 2020b). This work was conducted with authorisation from Ezemvelo KwaZulu-Natal Wildlife (Ordinary Permit: 4353/2018) and with clearance from the Stellenbosch University Research Ethics Committee (Protocol Number ACU-2019-9533). **Additional Information:** Supplementary Information is available online. Requests for additional materials should be addressed to CW. **Authors’ contributions:** CW and JM conceived and designed the project. CW, NPM and JM collected the data. CW led and performed the statistical analysis. CW led the writing of the manuscript, and all authors contributed to, and approved, the final manuscript.

## Abstract

Gut microbial communities regulate host physiology and health of humans and laboratory animals. The functional significance of these collective bacterial genomes (i.e. the microbiome) to the adaptive potential of wildlife hosts is still unknown. Studies demonstrating convincing examples of microbial flexibility to environmental change so far lack the experimental approaches to demonstrate the effect on host physiology. Invasive species provide natural experiments to tease apart these host-microbe relationships. However, no studies have investigated how microbial symbionts might mediate responses of invasive hosts’ physiology to environmental change. In this study, we examine whether invasive gut microbiomes have significantly diverged in their ability to respond to novel environmental change (i.e. a dietary challenge) compared to native gut microbiomes by performing reciprocal faecal microbial transplant (FMT) experiments in native and invasive guttural toad (*Sclerophrys gutturalis*) populations. Subsequently, we determine how the microbiome regulates host physiological changes in response to a dietary challenge. We show that invasive gut microbiomes exhibit higher microbial compositional and predicted functional flexibility to novel dietary change, compared to native gut microbiomes. This increased microbial flexibility is coupled with significant flexibility in energy harvesting. Furthermore, our results indicate that overall invasive gut microbiomes significantly upregulate energy harvesting and physiological performance of hosts, compared to native microbiomes. Our study is the first identifying gut microbiota as the sole factor contributing to the adaptive physiology of a vertebrate using a unique study design. These findings provide novel insights into the key role of gut microbial symbionts in increasing the invasive potential of its vertebrate host.

## INTRODUCTION

The vertebrate digestive system is host to diverse and complex microbial communities that play a fundamental role in the development, physiology and wellbeing of their hosts (O’Hara & Shanahan, 2006; Robinson et al., 2010; Cho & Blaser, 2012; Sommer & Bäckhed, 2013; Kohl & Carey, 2016; McKenney et al., 2018). Of all the abiotic and biotic factors that affect the composition of symbiotic microbial communities (reviewed in Costello et al., 2012 and Dickey et al., 2020), diet has the largest known impact on individual gut microbiomes (Turnbaugh et al., 2009; Faith et al., 2011; David et al., 2014; Kohl et al., 2014; Wilson et al., 2020). This is due to strong selection on gut microbial communities for their ability to degrade specific dietary molecules (Turnbaugh et al., 2009; David et al., 2014; Kohl et al., 2014). Dietary changes can, therefore, produce alterations in the host microbiota’s ability to mediate host processes such as digestion and energy acquisition (Bäckhed et al., 2005; Turnbaugh et al., 2006; De Angelis et al., 2020), vitamin synthesis (Zmora et al., 2019), immunomodulation (Longman & Littman, 2015; Leigh & Morris, 2020; De Angelis et al., 2020), pathogen defence (Dethlefsen et al., 2007; Longman & Littman, 2015) and possibly host physiology (Alberdi et al., 2016; Fontaine & Kohl, 2020). Given the dynamic association between diet and the gut microbiome, inferring how microbiota respond to dietary changes and subsequently, shifts in hosts’ physiology presents a meaningful challenge.

By using an ecological approach, the gut can be viewed as a distinct microbial habitat where gut microbial communities are affected by similar processes as those explained by island biogeography theory (i.e. immigration, colonisation, and extinction) and community ecology theory (i.e. deterministic, neutral, and historic processes of community assembly) (Walter & Ley, 2011; Costello et al., 2012; Delong, 2014; Bletz et al., 2016). Within this framework, entering a new environment by an alien host is likely to involve dietary changes, and thus, a host’s gut microbial community can be predicted to adjust in response to these changing ecological conditions (i.e. microbial flexibility). However, microbial communities, like other ecological communities, can also display “resistance” to dietary (or environmental) change, i.e. communities remain essentially unchanged despite disturbance (Moya & Ferrer, 2016). Microbial resistance to diet alterations has been demonstrated in human and laboratory animal populations and is usually coupled with an inability to maintain microbial function, and host health and physiology (Smith et al., 2013; Jandhyala et al., 2015; Sonnenburg & Bäckhed, 2016). In contrast, a few studies concerning wildlife populations show that while animals display microbial flexibility in response to disturbance, this flexibility is coupled by functional maintenance or redundancy (Bletz et al., 2016; Voolstra & Ziegler, 2020; Webster et al., 2020; Fontaine & Kohl, 2020). Studying wildlife populations and their microbial symbionts in natural conditions are, therefore, of great importance in order to elucidate the natural responses of microbial communities and their hosts to environmental change.

If microbial flexibility in response to novel environmental change is coupled with microbial functional redundancy, then maintenance of host physiology, health and/or fitness would be expected. However, no studies have demonstrated conclusively that gut microbial responses to changing host environments lead to measurable effects on host health or physiology in fully natural systems (Hauffe & Barelli, 2019; Greyson-Gaito et al., 2020, but see Bletz et al., 2016; van Opstal & Bordenstein, 2019; Gomes et al., 2020; Greenspan et al., 2020). Research on humans and laboratory animals has shown that the gut microbiome acts as an important mediator of the relationship between dietary change and host physiology (Turnbaugh et al., 2006; Sonnenburg & Bäckhed, 2016). To infer direct impacts of the gut microbiome on host physiology and health, laboratory studies implement faecal microbial transplants (FMT).

However, laboratory studies are unable to account for natural environmental variability experienced by the host and its associated gut microbiome (Greyson-Gaito et al., 2020). Moreover, laboratory animals normally have significantly different microbiomes compared to their wild counterparts, which makes it difficult to tease apart the ecology and evolution of wild host-microbial associations (Dethlefsen et al., 2007). To understand the processes impacting natural microbial variation and how the microbiome mediates host physiological responses in changing environments, we need to move beyond laboratory systems and expand to studying microbiomes in wild populations.

Biological invasions provide a valuable opportunity as natural experiments to investigate evolutionary responses of populations and their microbial symbionts to changing environmental conditions. Dietary and intestinal flexibility of introduced populations have been shown to contribute greatly to their success in novel environments (Caut et al., 2008; Banks et al., 2008; Kidera et al., 2008; Ruffino et al., 2011; Redford et al., 2012). The guttural toad (*Sclerophrys gutturalis*) is an invasive amphibian introduced ~20 years ago into periurban Cape Town, from its known source population in Durban, South Africa (Telford et al., 2019). Previously, massive parallel sequencing of invasive guttural toad population gut microbiomes showed that the Cape Town toad population gut microbiome has diverged to become compositionally, phylogenetically and functionally distinct from its source population (Wagener et al., *in review*). Furthermore, toads within the Cape Town invasion exhibit increased endurance under dehydrated conditions (Vimercati et al., 2018), sustained resource investments to growth (Vimercati et al., 2019), and behavioural shifts to conserve water (Madelaire et al., 2020). Guttural toads in these areas provide us with a well-studied system to investigate whether gut microbial communities have diverged in their flexibility to respond to changing environmental conditions and how this change impacts predicted microbial functionality and host physiology.

We used this amphibian host system to test (1) the hypothesis that invasive microbiomes will exhibit a higher degree of microbial flexibility or plasticity following environmental change (in this case, a novel dietary challenge) and (2) if a higher degree of microbial flexibility (or lack thereof) will allow the maintenance of similar predicted microbial functionality and host physiology (in terms of resource intake and physiological performance) in response to environmental change. The reciprocal transfer of individuals and/or their microbes between different habitats is a classical technique used in evolution and invasion ecology to ascertain how and by what mechanisms individuals or their microbes adjust to environmental change. Therefore, we used this approach combined with faecal microbial transplants (FMTs) to reciprocally transfer invasive and native gut microbial communities to individual hosts in Durban (native population) and Cape Town (invasive population) in order to answer these hypotheses.

## MATERIALS AND METHODS

### Study sites

Adult female guttural toads (*Sclerophrys gutturalis*) were collected in their native range (Durban, 29° 83′ S, 30° 93′ E; 29 May to 10 June 2019) and invasive range (Cape Town, 33° 99′ S, 18° 44′ E; 2 to 30 March 2019). We captured animals of a single sex (females) as intersex microbiome differences may confound results. A total of 48 individuals were collected for experimental trials in each area. A further 34 individuals were collected in each area to serve as faecal material donors.

### Collection and preparation of donor faecal material

Within each sampling area, adult female toads were captured by hand after sunset (19:00 h). Immediately after capture, toads were weighed (to the nearest 0.01 g; WTB 2000, Radwag, Radom, Poland), and their snout-to-vent length (SVL) was measured using a digital calliper (to the nearest 0.01 mm, Mitutoyo). Toad sex was confirmed through visual inspection for white colouration of the gular region and a greater than 40 mm SVL measurement (Baxter-Gilbert et al., 2020). Female toads were then placed individually in plastic containers (195 x 195 x 180 mm), which had been sterilized with a 10% bleach solution followed by 70% ethanol. Fresh faecal matter was processed and stored within 8 hours of defecation. After 8 hours of capture, native toads were released, whereas invasive toads were euthanized by immersion in a 1 gl^−1^ solution of tricaine ethane sulfonate (MS-222) for 20 minutes according to legal requirements (Davies et al 2020b).

At least 0.3 g faecal material from each donor was suspended within a 1.0 mL sterile saline solution (0.9% NaCl). Glycerol (90.08%) was then added to obtain a final concentration of 10% (Satokari et al., 2015). Glycerol is used as a cryoprotective agent for frozen faecal samples (Gough et al., 2011; Hamilton et al., 2012) and ensures microbial viability for up to 6 months in −80 °C (Costello et al., 2015). The final suspension was clearly labelled and stored at −80°C.

### Recipient Toad Housing

Recipient toads were housed in mesocosm enclosures created from plastic pools (310 L, 1.98 m in diameter and 0.38 m deep. Two enclosures were placed within each sampling area, Durban and Cape Town (Figure 1). Use of outdoor mesocosms increases environmental relevance and maximizes the benefits of field experiments by maintaining relatively controlled environments while incorporating natural elements, such as daily variation in temperature and rainfall (Rowe & Dunson, 1994). Screen mesh was placed on top of mesocosms to prevent toads from escaping and predators from entering the enclosures. Blacklights were suspended 30 cm aboveground inside each enclosure and illuminated each night to attract insects. Toads were, therefore, sustained on a ‘natural diet’. Every second day fresh soil and leaf litter was placed inside enclosures.

**Figure 1.**
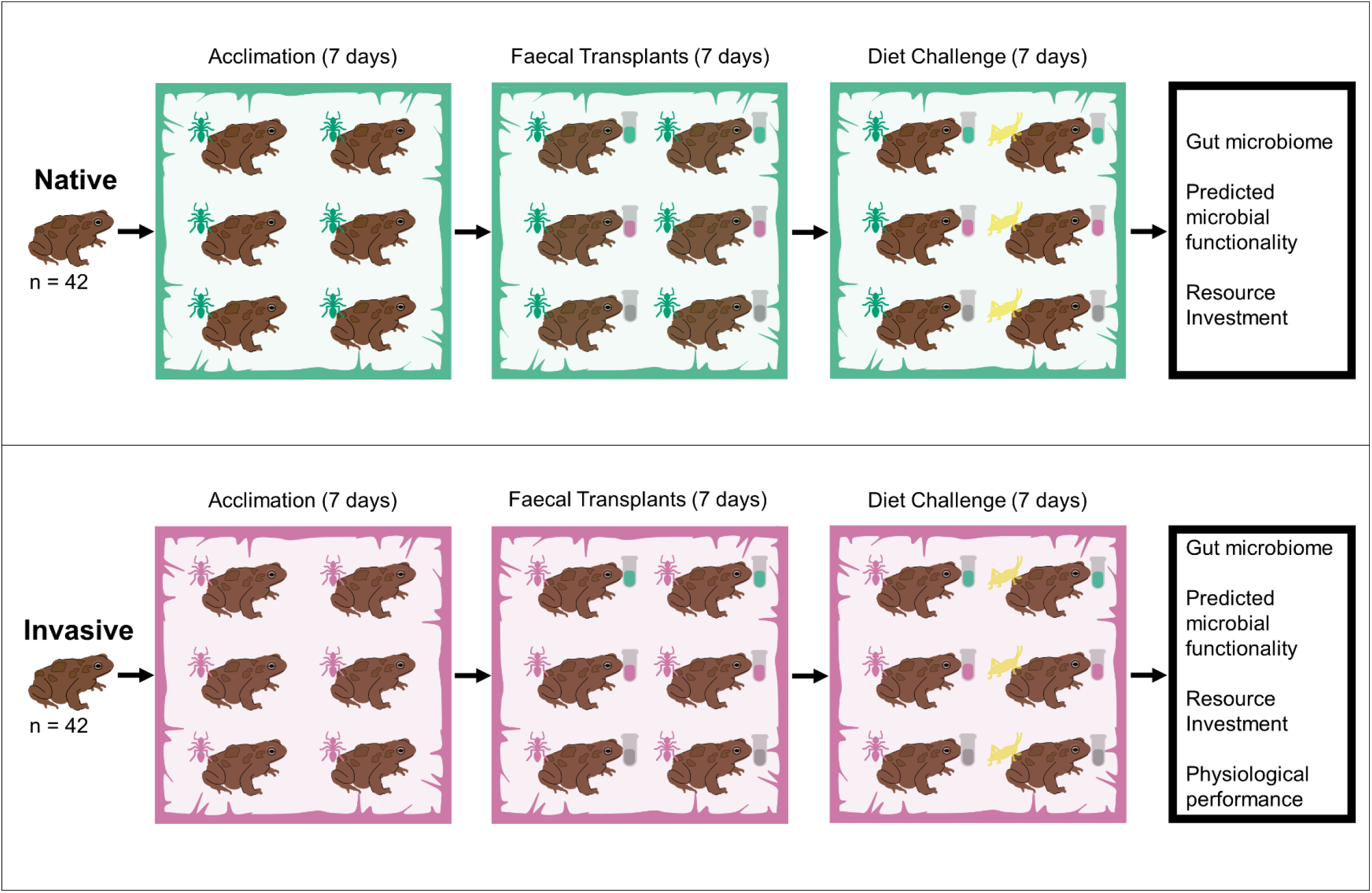
Overview of experimental design. Experiments were completed on native and invasive guttural toads (*Sclerophrys gutturalis*) in their native, Durban (blue), and invasive, Cape Town (pink), respectively. Toads were captured in the respective regions and allowed to acclimate to mesocosms for seven days while sustained on a natural diet (blue or pink ants). After seven days of acclimation to mesocosms, guttural toads were colonized with the gut microbiome of native toads (blue vials, of Durban origin) and invasive toads (pink vials, of Cape Town origin). Additionally, a final group of toads acted as a control (grey vials). This created six FMT (faecal microbial transplant) treatment groups across experimental areas: Cape Town control, Cape Town self-transplant, Cape Town transplant, Durban control, Durban self-transplant and Durban transplant. After seven days, toads were then subjected to one of either two diets: a continuation of their natural diets (blue or pink ants) or a novel dietary challenge (yellow crickets), while the FMT treatments continued for each group as before. Gut microbial composition, predicted functional capabilities, body mass, lean structural mass, body fat % and liver mass of all toads was measured after experimental trials. Endurance and speed were additionally measured in the invasive region.

Before placement in enclosures, toads’ snout-to-vent length and mass were measured. Individuals were tagged using 8 mm PIT tags, which are small glass capsules with an electromagnetic coil (Guimaraes et al., 2014). The tag was placed in a 15-gauge hypodermic needle and injected underneath the skin above the dorsal lymph sac (following Donnelly et al., 1994). Afterwards, toads were randomly assigned to each of the two enclosures in each area. Toads were then allowed seven days to acclimate to enclosure conditions (Figure 1). Throughout the acclimation and experimental periods, toads’ body mass was measured daily.

### Faecal Microbial Transplants

After the acclimation period, toads were administered one of three faecal microbial transplant (FMT) treatments: control, invasive or native, for seven days (Figure 1). Toads on the control treatments received a 10% glycerol solution dissolved in 1.0 mL sterile saline (0.9% NaCl). Toads under the invasive treatment received approximately 0.3 g invasive toad faecal samples (of Cape Town origin) suspended in a 1.1 mL 10% glycerol and saline solution. Toads under the native treatment received approximately 0.3 g native toad faecal samples (of Durban origin) suspended in a 1.1 mL 10% glycerol and saline solution.

Each day, faecal suspensions were thawed to room temperature. Sterile saline solution (0.9% NaCl) was added to each aliquot to obtain the desired suspension volume. Solutions were fed to toads via a neonatal feeder. These FMT treatments were repeated in each sampling area, Durban (native area) and Cape Town (invasive area), creating six FMT treatment groups across the sampling areas: Durban control, Durban self-transplant (i.e. Durban native toads fed native faecal material), Durban transplant (i.e. Durban native toads fed invasive faecal material), Cape Town control, Cape Town self-transplant (i.e. Cape Town invasive toads fed invasive faecal material), Cape Town transplant (i.e. Cape Town invasive toads fed native faecal material) (Figure 1).

### Diet Challenge

After the initial seven days of FMT supplementation to natural diets in mesocosms, toads in each FMT treatment group were exposed to one of two diets along with their normal supplementation of gut samples (Figure 1). Toads were either continued on a natural diet or were exposed to a novel diet challenge. Toads exposed to the dietary challenge were fed four large house crickets (*Acheta domesticus* (Linnaeus)) every second day. The dietary changes were continued for seven days.

### Faecal Sample Collection

After completion of experiments, faecal samples were collected from all individuals. As before, toads were placed individually in sterilized plastic holding containers (195 x 195 x 180 mm) and at least 0.3 g faecal matter was obtained from each individual. Samples were immersed in 1.0 *RNAlater*^TM^ within 2 ml sterile polypropylene tubes (Ambion, Austin, TX). After approximately 48 hours, faecal samples were centrifuged (2 min at 10 000 x g), the supernatant was removed, and the pellet stored at −80 °C. Empty tubes containing *RNAlater*^TM^ and glycerol solutions were kept as negative controls for DNA processing.

### DNA Extraction and purification

The DNeasy^®^ PowerSoil^®^ kit (QIAGEN, Hilden, Germany) was used, according to the manufacturer’s protocol, to extract metagenomic DNA from 0.25 g of each faecal sample. DNA extracts were stored at −80 °C until further processing. No template and template from blank filters were included as negative controls throughout the entire process from DNA extraction to PCR amplification.

DNA samples were quantified using the Qubit 4.0 Fluorometer (ThermoFisher Scientific) and the Qubit 1x dsDNA HS assay kit (ThermoFisher Scientific) according to the manufacturer’s protocol. To determine the purity of the metagenomic DNA samples, spectrophotometry was performed on the NanoDrop^®^ ND-1000 (ThermoFisher Scientific). Genomic quality scores (GQS) were determined on the LabChip GXII Touch using the DNA Extended Range LabChip and Genimic DNA Reagent Kit (PerkinElmer, Waltham, MA, USA), according to the manufacturer’s protocol.

### PCR amplification

The V3 and V4 hypervariable regions of the rRNA were targeted during sequencing. Target 16S rRNA sequences were amplified using the universal bacterial primer set, 314F 5’ – CCTACGGGNGGCWGCAG – 3’ and 785R 5’ – GACTACHVGGGTATCTAATCC – 3’ (Klindworth et al., 2013). Fragments were amplified from 5 ng metagenomic DNA in a reaction volume of 20 µl (0.5 µM of each primer, 200 µM dNTPs, 0.4 U Phusion hot-start II high-fidelity (HF) DNA polymerase and 1 x Phusion HF buffer) with a final concentration of 1.5 mM MgCl_2_. Polymerase chain reactions (PCRs) were performed on the SimpliAmp^TM^ Thermal Cycler (ThermoFisher Scientific). Initial template DNA denaturation at 98 °C for 30 sec was followed by 25 cycles consisting of 98 °C for 10 sec, 58 °C for 30 sec and 72 °C for 30 sec; with a final product extension at 72 °C for 10 min.

Presence of amplified products were verified on the PerkinElmer LabChip^®^ GXII Touch (PerkinElmer, Waltham, MA, USA), using the X-mark chip and HT DNA NGS 3K reagent kit, according to the manufacturer’s protocol. PCR products were then purified with 1.8x volume Agencourt™ AMPure™ XP reagent (Beckman Coulter, Brea, CA, USA) and eluted in 25 µl nuclease-free water. Purified amplicons were quantified on the Qubit 4.0 Fluorometer using the Qubit 1x dsDNA HS assay kit (ThermoFisher Scientific), according to the manufacturer’s protocol.

### Library Preparation

Library preparation from 100 ng PCR product per sample was performed using the NEXTflex DNA Sequencing Kit (Bio Scientific Corporation) according to the manufacturer’s protocol. Approximately 40 µl from each purified PCR product was end-repaired at 22 °C for 30 min using 3 µl End-repair enzyme mix and 7 µl End-repair buffer in a final volume of 50 µl. The end-repaired products were purified with 1.8x volume Agencourt™ AMPure™ XP reagent (Beckman Coulter). About 19 µl purified, end-repaired product was ligated to 4 µl IonCode™ Barcode Adapter (ThermoFisher Scientific) with the addition of 31.5 µl Ligation mix at 22 °C for 15 min. The adapted-ligated, barcoded libraries were then purified with 1.8x Agencourt™ AMPure™ XP reagent (Beckman Coulter) and quantified using the Ion TaqMan Library Quantitation Kit (ThermoFisher Scientific). Using the StepOnePlus™ Real-time PCR system (ThermoFisher Scientific), qPCR amplification was performed. Library fragment size distributions were assessed on the LabChip^®^ GXII Touch (PerkinElmer, Waltham, MA, USA), using the X-mark chip and HT DNA NGS 3L reagent kit according to the manufacturer’s protocol.

### Sequencing

Massive parallel sequencing was performed on the Ion GeneStudio™ S5 Prime System using the Ion S5™ Sequencing solutions and reagents according to the manufacturer’s protocol.

### Sequencing Data Pre-processing

Resulting sequences were stored in FASTQ formatted files generated for each sample. Single-end raw reads (11 865 157) were imported into QIIME2 (version 2020.2) for pre-processing (Bolyen et al., 2019). The divisive amplicon denoising algorithm (DADA2) plugin was used to de-noise sequencing reads (Callahan et al., 2016). Briefly, low-quality sequences (sequences < 400 bp in length and < 20 quality score, sequences containing ambiguous characters, unreadable barcodes or without primer sequences), chimeric sequences and singletons were removed using default DADA2 parameters. The resulting sequences were then used to generate amplicon sequence variants (ASVs) for downstream analyses. This resulted in 6 973 959 sequences ranging from 53296 to 154 822 sequences per sample representing a total of 13 986 unique ASVs. ASV sequences were aligned with mafft (Katoh & Standley, 2013; q2-alignment plugin), high entropy positions were filtered from the resulting alignment (Lane et al., 1991), an unrooted tree was constructed with FastTree (Price et al., 2010; q2-phylogeny plugin) and the tree was rooted using midpoint rooting. Taxonomy was assigned to ASVs with a classify-sklearn classifier trained against the most recent SILVA 16S rRNA gene reference database (release 138) (Quast et al., 2013; q2-feature-classifier plugin). The ASV table, phylogenetic tree and assigned taxonomy table was used in all downstream analyses.

The ASV table and its corresponding phylogenetic tree was additionally used to predict functional profiles of samples through the PICRUSt2 (Phylogenetic Investigation of Communities by Reconstruction of Unobserved States 2) pipeline in QIIME2 (Douglas et al., 2020; q2-picrust2 plugin) and the KO Database of Molecular Functions by ortholog annotation (KEGG orthologues, KO, https://www.genome.jp/kegg/ko.html).

All negative controls were removed due to low sequence number (< 100) and sequence quality score (< 20). Removal of contaminant sequences were, therefore, not required.

### Performance

The effect of diet and FMT treatment on performance was tested in 44 toads from Cape Town (no Durban toads were used). Toads were tested on an indoor circular racetrack (4.1 m) using a rubber grip mat as a substrate (Vimercati et al., 2018). Performance trials were completed during the day between 09h00 – 18h00. Each toad was individually placed on the racetrack and stimulated to hop by gently tapping it on the urostyle with a brush. To standardize tapping time, toads were tapped by a single operator (CW) at intervals of 1 s after each hop. For each toad, we counted the number of laps and therefore the distance (4.1 m per lap) moved until it did not voluntarily hop for 10 consecutive taps (i.e. exhaustion). For each lap around the racetrack we also recorded the time taken until exhaustion.

### Dissections

After faecal sample and performance data collection, toads from Cape Town (n = 48) and Durban (n = 48) were euthanized by immersion in a 1 gL^−1^ solution of tricaine ethane sulfonate (MS-222) for 20 minutes. The carcasses were frozen (−20° C) in labelled plastic bags until dissection. In the laboratory, after defrosting each specimen at ambient temperature, individuals were weighed (± 0.01 g) and their SVL was measured using digital callipers (± 0.01 mm). Fat bodies and liver were weighed after removing each organ (± 0.01 g). Tissues were patted dry with a paper towel before weighing. The percentage of body mass composed of fat reserves (hereafter, body fat %) was obtained from the ratio between the mass of fat bodies and body mass (Brown et al., 2011). Lastly, individuals were fully eviscerated and weighed to obtain lean structural mass (± 0.01 g).

### Statistical analysis

Preliminary analyses showed that body mass was positively correlated with SVL. Therefore, the body condition (or scaled body mass) was calculated following Peig and Green (2009) and Vimercati et al. (2019). Body condition of toads was used as a covariate in all downstream microbiome analyses. Preliminary analyses indicated that faecal microbial transplants were successful as there were no significant differences of microbial composition between control and self-transplant groups within each experimental area (Table S1).

All statistical analyses were performed in R version 3.6.2 (R Core Team, 2019). Metadata, ASV table, taxonomy and phylogenetic tree was imported using the qiime2R package (v0.99.13, Bisanz, 2018). A phyloseq object was built from these datasets using the phyloseq package (v1.30.0, McMurdie & Holmes, 2013). Prior to all downstream analyses alpha rarefaction curves were inspected to assess sequencing depth (figure S3.1). Visual inspection confirmed that sequencing depth was adequate for each sample with regards to number of ASVs detected. ASV counts of each sample were then filtered, removing ASVs present in less than 5% of the samples, and according to the read depth of each sample using the phyloseq and microbiomeutilities packages (v0.99.02, Shetty & Lahti, 2018).

Diversity metrics inverse Shannon diversity, Evenness, Chao1 species richness and Faith’s phylogenetic diversity metrics was calculated using the vegan package in R (v2.5.6, Oksanen et al., 2007). The inverse Shannon diversity metric incorporates both measures of species richness and abundance. Evenness estimates how similar in abundance species in a sample are, while Chao1 estimates the asymptote on a species accumulation curve to determine species richness. Faiths’ phylogenetic diversity metric measures the cumulative branch lengths from randomly sampled species from each sample. Generalized linear models (GLM) were used to determine the effect of FMT treatment, dietary change, their interaction and body condition (dependent variables) on alpha diversity metrics (response variables). Prior to analyses, model assumptions (e.g. normality, homogeneity, and independence) were assessed. None of the diversity estimates, except phylogenetic diversity within the Durban area group, met model assumptions of normality. To meet model assumptions, inverse Shannon and phylogenetic diversity metrics were log-transformed and Chao1 and Pielou diversity estimates were 1/x transformed for the Cape Town group. For the Durban group, inverse Shannon and Chao1 estimates were square-root transformed and Pielou diversity estimates were 1/x transformed. Relative variable importance of competing models was evaluated using Akaike information criterion (AIC). Chi-square values and associated p-values were investigated to examine the effect of the response variables on the dependent variables.

The interaction effect of FMT treatment and dietary change on microbiome CLR- and PHILR-composition matrices was examined using PERMANOVA analyses. CLR- and PHILR-metrics are equivalent to the Bray-Curtis and Unifrac beta diversity metrics, but account for the compositional nature of the data (Gloor et al., 2017). Feature tables containing read counts were first subjected to centre log-ratio (CLR)- and PHILR-transformation using phyloseq and philr packages (v1.12.0, Silverman et al., 2017). Euclidean distance matrices were constructed from the transformed ASV count tables through the adonis function (vegan package). Distance matrices were then subjected to PERMANOVA analyses (999 permutations) to evaluate the effect of FMT treatment, diet change, their interaction and body condition on toad gut microbial composition. As PERMANOVA is sensitive to differences in dispersion of data within groups (assumes a homogenous within-group dispersion), we inspected this assumption with the betadisper and permutest functions of vegan. CLR- and PHILR-transformed Euclidean distance matrices were also used in principle component analyses (PCoA) to visualize the responses of population gut microbial communities to novel dietary changes.

To investigate differential abundance of ASVs, likelihood ratio tests (LTR) were employed through the DeSeq2 package (v1.26.0, Love et al., 2014). This test was implemented using a full model with body condition and the interaction effect of FMT treatment and diet change against a reduced model with body condition as the only predictive variable. Prior to analyses, read counts were normalized using a regularized logarithm. The Benjamini-Hochberg method for reducing false discovery rate (FDR) was employed with a cut-off of < 0.05 for identifying differentially abundant microbes. Corresponding log-fold change, p-values and FDR-adjusted p-values were estimated. To investigate differences in abundances of ASVs between FMT treatments and diet changes, pairwise comparisons were also performed using DeSeq2.

Functional components of bacterial communities were assessed. Prior to all analyses pathway abundances derived from the PICRUSt2 pipeline were filtered for pathways with > 5 counts. Data was then subjected to compositional (beta diversity) and differential abundance analyses similar to those described above.

Body mass, lean structural mass, and liver mass were positively correlated with SVL and thus, we calculated the scaled mass index for these variables following Pieg and Green (2009) and Vimercati et al. (2019). To determine the effect of FMT treatment, diet change, and their interaction on the scaled body mass, lean structural mass, body fat % and scaled liver mass (response variables) of toads in each area (Cape Town and Durban), generalized linear models (GLM) were employed as described above. Prior to analyses, model assumptions (e.g. normality, homogeneity and independence) were assessed. Body mass and lean structural mass was log-transformed and scaled liver mass and body fat % was square-root transformed in order to meet model assumptions of normality.

Finally, mixed effects models (GLMM) were used to determine the effect of FMT treatment, diet change, and their interaction on performance measures; endurance (m) and speed (m.s^−1^). Snout-to-vent length (SVL) of toads was included as a covariate in analyses and trial number as random factor. Both performance measures were square-root transformed in order to meet model assumptions of normality. Relative variable importance of competing models was evaluated using Akaike information criterion (AIC). To evaluate the variance of data explained by each model, marginal (fixed effects) and conditional (fixed and random effects) R2 was calculated according to Nakagawa and Schielzeth (2013) using the ‘r.squaredGLMM’ function in the package MuMIn (v1.43.15, Barton, 2009). F-statistics and associated p-values were investigated to examine the effect of fixed effects on the dependent variables.

## RESULTS

### Invasive gut microbiomes display microbial flexibility in response to a dietary challenge

The alpha diversity of gut bacterial communities does not differ across toad microbiomes or in response to dietary change (Figure S1 and Figure S2; Table S2). However, different toad gut microbiomes display varying responses (i.e. microbial flexibility) to a novel dietary challenge (significant interaction effect; Cape Town: PERMANOVA, Pseudo-F_(1, 41)_ = 1.33, *p* < 0.001 and Durban: PERMANOVA, Pseudo-F_(1, 41)_ = 1.27, *p* < 0.05, Table 1, Figure 2A; Figure 2B). Toads with invasive gut microbiomes (i.e. Cape Town control, Cape Town self-transplant, and Durban transplant) significantly shift their gut microbial composition in response to a dietary challenge (PERMANOVA *p*, < 0.01 for all comparisons, Table S1). Toads with native gut microbiomes (i.e. Durban control, Durban self-transplant, and Cape Town transplant), on the other hand, display microbial resistance towards a dietary challenge, with no significant differences observed between toads with native microbiomes on natural or novel diets (PERMANOVA, *p* > 0.05 for all comparisons, Table S1). Responses of invasive toads’ microbiomes to a dietary challenge is not the result of dispersion variation between microbiomes (BETADISPR, *p* > 0.05, Figure 2C; Figure 2D). Body condition also has no effect on the gut microbial shifts observed in guttural toad hosts (Table S3). Similar patterns are observed for phylogenetic diversity (Table S1, Table S3, Figure S3).

**Figure 2.**
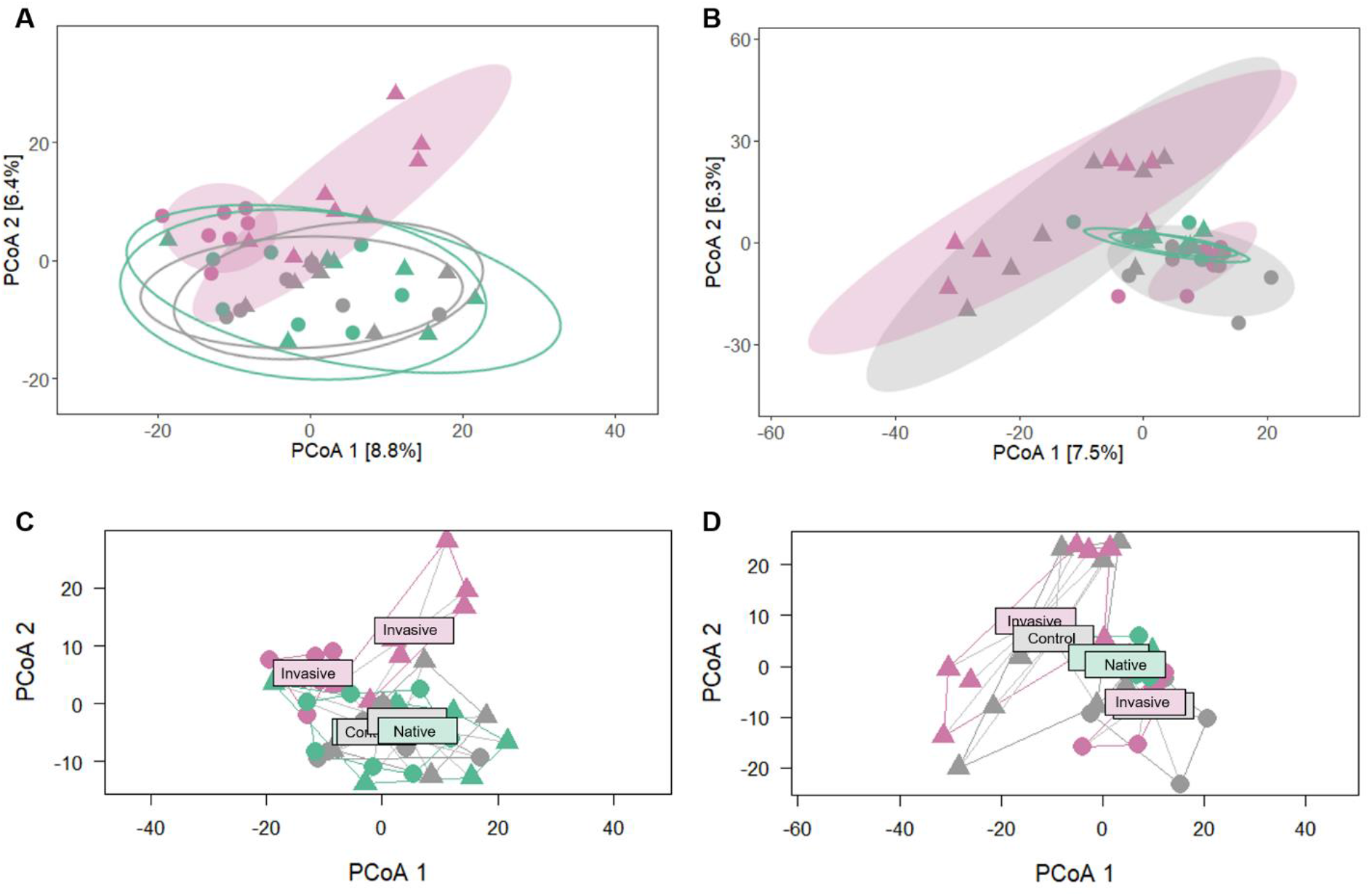
Principle Coordinates Analysis (PCoA) of CLR-Euclidean compositional beta diversity demonstrating responses of guttural toads (*Sclerophrys gutturalis*) colonized by native (blue), invasive (pink), and control (glycerol, grey) toad gut microbial communities to natural diets (circles) and a novel dietary challenge (triangles). Experiments were completed in the toads’ native range, Durban (A, C) and invasive range Cape Town, South Africa (B, D). PERMANOVA tests indicated that invasive gut microbiomes (shaded ellipses: Cape Town self-transplant, Cape Town control and Durban transplant) significantly shift their gut microbial composition in response to a dietary challenge, whilst native gut microbial communities (empty ellipses: Durban self-transplant, Durban control and Cape Town transplant) show no response. Permutational test of dispersions (PERDISP) showed responses were not the result of variation in dispersion.

Of the 732 (Cape Town) and 539 (Durban) differentially abundant ASVs, more ASVs present in the invasive gut microbiome (i.e. Cape Town control, Cape Town self-transplant and Durban transplant) become differentially abundant in response to a dietary challenge, compared to those present in the native gut microbiome (i.e. Durban control, Durban self-transplant, and Cape Town transplant) (Table S4). The invasive gut microbiome significantly alters microbial abundance of 395 ASVs (Cape Town self-transplant) and 302 ASVs (Cape Town control) in response to a dietary challenge in Cape Town (invasive area) (Table S5). Only 185 ASVs (Cape Town transplant) of the native gut microbiome shift in response to a dietary challenge (Table S5). In Durban (native area), the invasive gut microbiome significantly alters microbial abundance of 174 ASVs (Durban transplant) while 162 ASVs (Durban self-transplant) and 113 ASVs (Durban control) of native gut microbiomes shifts in response to a dietary challenge (Table S5).

### Microbial flexibility in invasive toads is coupled with predicted functional flexibility

Guttural toads’ predicted microbial functional diversity is determined by both the microbiome (FMT treatments) and diet (Table S6, Figure S3C; Figure S3D). However, significant interaction effects are only present in hosts from Cape Town (Cape Town: PERMANOVA, Pseudo-F_(1, 41)_ = 1.57, *p* < 0.05 and Durban: PERMANOVA, Pseudo-F_(1, 41)_ = 1.01, *p* > 0.05; Table S6). Invasive gut microbiomes (i.e. Cape Town control, Cape Town self-transplant and Durban transplant) significantly shift their predicted gut microbial functional capabilities in response to a dietary challenge (PERMANOVA, *p* < 0.01 for all comparisons, Table S7). Toads with native gut microbiomes (i.e. Durban control, Durban self-transplant and Cape Town transplant), on the other hand, display no functional variation in response to a dietary challenge (PERMANOVA, *p* > 0.05 for all comparisons, Table S7). Changes of predicted functional capabilities is not the result of dispersion variation (BETADISPR, *p* > 0.05, Figure S3G; Figure S3H). Body condition also has no effect on the microbial functional differences (Table S6).

In Cape Town and Durban, 53 and 158 functional pathways, respectively, are differentially abundant across FMT treatments and diets (Table S8). The invasive gut microbiome significantly alters the abundance of 36 (Cape Town self-transplant) and 21 (Cape Town control) functional pathways in response to a dietary challenge in Cape Town (table S9). Only 11 (Cape Town transplant) functional pathways of the native gut microbiome shift in response to dietary challenge. In Durban, the invasive gut microbiome significantly alters the abundance of 48 (Durban transplant) functional pathways while 35 (Durban self-transplant) and 46 (Durban control) functional pathways of the native gut microbiomes shift abundance in response to novel dietary challenge (table S9).

### Invasive gut microbiomes stimulate increased resource intake

Scaled body mass and lean structural mass does not vary across toad FMT treatments or diets (Table S10; Figure S4). Body fat % and scaled liver mass, on the other hand, does vary in response to a novel dietary challenge depending on the FMT treatment (or gut microbiome) of toads (significant interactions effects; body fat %: GLM, *p* < 0.05 in both experimental areas; and scaled liver mass: GLM, *p* < 0.05 in both experimental areas; Figure 3; Table S10). Toads with invasive gut microbiomes (i.e. Cape Town control, Cape Town self-transplant and Durban transplant) have significantly higher body fat % and scaled liver mass when subjected to a dietary challenge, while toads with native gut microbiomes (i.e. Durban control, Durban self-transplant and Cape Town transplant) show no response (Table S11). Additionally, toads with invasive gut microbiomes have a significantly higher body fat % and scaled liver mass than toads with native gut microbiomes, irrespective of diet (Table S12). Pairwise comparisons also show that there were no differences of body fat % and scaled liver mass between toads in their respective control and self-transplant groups (Table S11).

**Figure 3.**
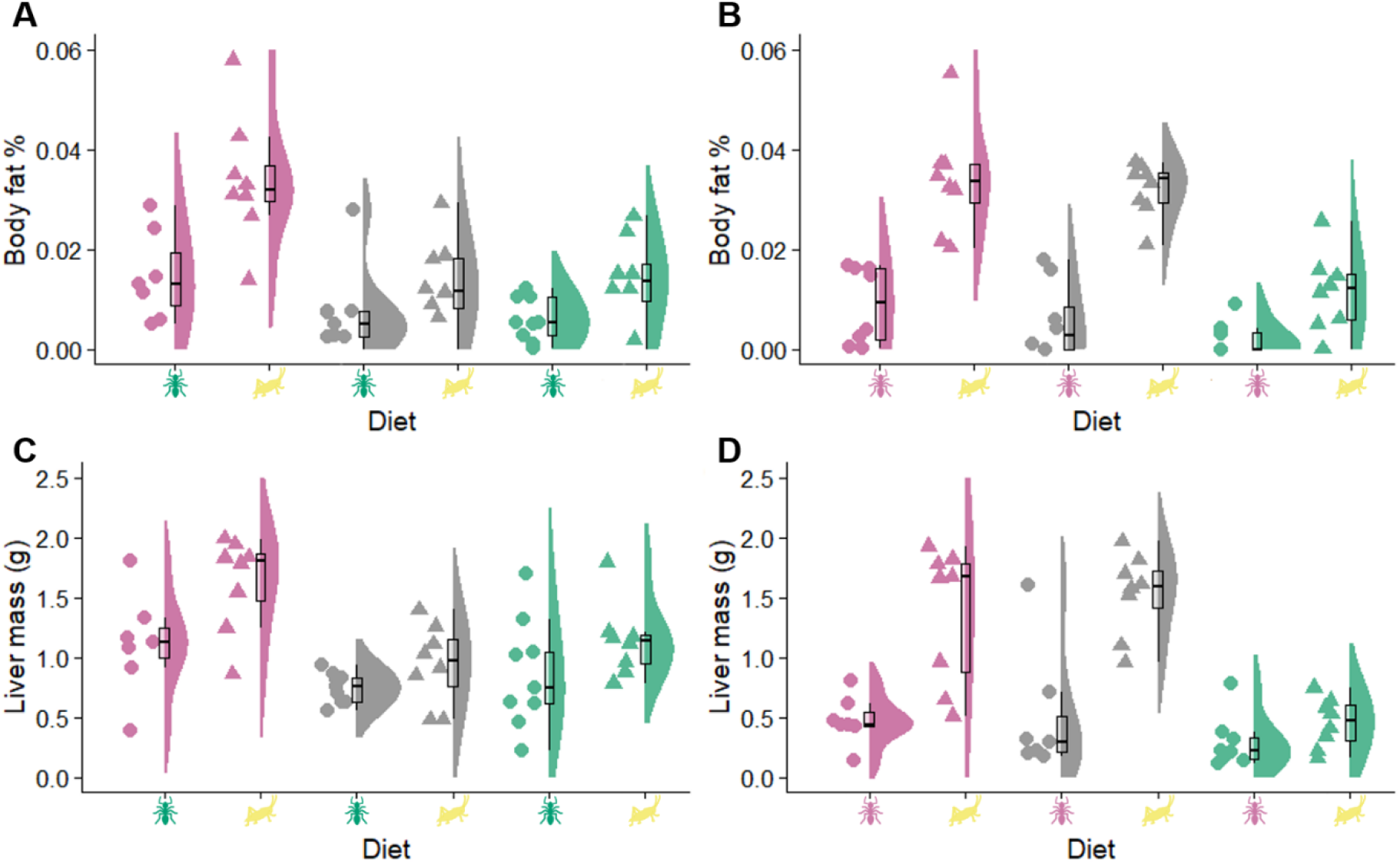
Body fat % and liver mass of guttural toads (*Sclerophrys gutturalis*) colonized by native (blue), control (glycerol, grey) and invasive (pink) toad gut microbial communities and subsequently subjected to two diets, natural (native blue and invasive pink ants) and novel dietary challenge (yellow crickets). Experiments were completed in the toads’ native range, Durban (A, C) and invasive range Cape Town, South Africa (B, D). Toads with invasive gut microbiomes significantly increase body fat % and liver mass when subjected to a dietary challenge, while toads with native gut microbiomes show no significant differences (GLM, *p* < 0.05). Toads with invasive gut microbiomes also has a higher overall body fat % and liver mass compared to toads with native gut microbiomes (GLM, *p* < 0.05).

### The gut microbiome alters physiological performance irrespective of dietary change

The gut microbiome impacts the distance travelled and speed of guttural toads, i.e. only FMT treatment has a significant effect on the performance of guttural toads (Figure 4; Table S13). Guttural toads with invasive gut microbiomes (i.e. Cape Town control and Cape Town self-transplant) attain significantly longer distances and higher performance speeds than those with native gut microbiomes (i.e. Cape Town transplant) (Table S14). There are no differences of performance between Cape Town control and self-transplant groups (Table S14).

**Figure 4.**
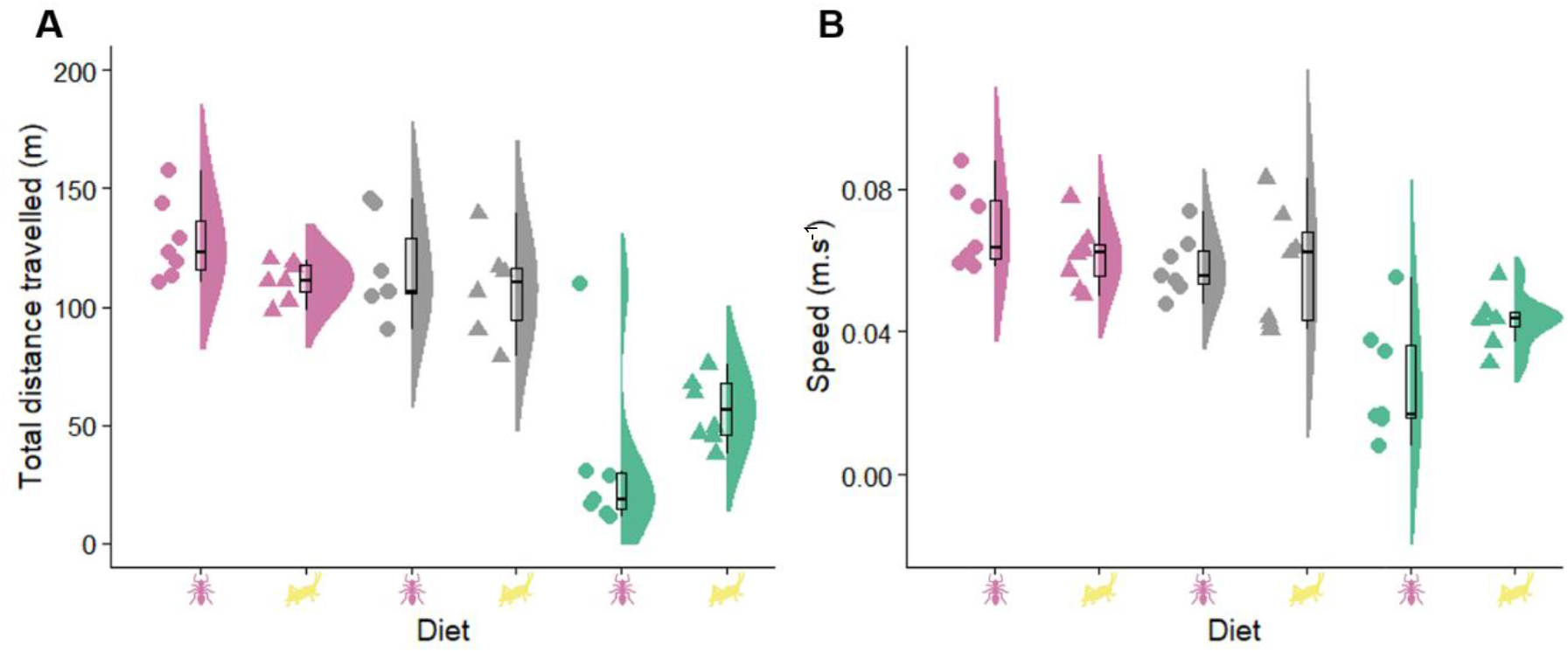
Physiological performance, total distance travelled (m) (A) and speed (m.s^−1^) (B), of guttural toads (*Sclerophrys gutturalis*) colonized by native (blue), control (glycerol, grey) and invasive (pink) toad gut microbial communities and subsequently subjected to two diets, natural (native blue and invasive pink ants) and novel dietary challenge (yellow crickets). Toads display no significant difference of physiological performance between diets (GLM, *p* > 0.05). However, toads with invasive gut microbiomes have significantly higher physiological performance compared to toads with native gut microbiomes (GLM, *p* < 0.05).

## DISCUSSION

How complex host-associated microbial communities respond to environmental change is of great interest in the fields of ecology and evolution. In this study, we demonstrate the varying adaptive responses of invasive and native gut microbial communities to novel diets. We show that the invasive *Sclerophrys gutturalis* gut microbiomes exhibit a higher degree of microbial flexibility enabling them to rapidly respond to novel dietary change compared to their native microbiomes. While there are similar studies that provide compelling evidence for microbial flexibility in wildlife populations (reviewed in Hauffe & Barelli, 2019), these case studies are few and lack experimental approaches and rigour required to determine that microbial flexibility impacts host fitness. Our results indicate that increased microbial flexibility of invasive gut microbiomes facilitates an increased flexibility of microbial predicted functional capabilities and resource investments or energy harvesting in toad hosts. Furthermore, we demonstrate that the invasive gut microbiome facilitates increases in resource investment (i.e. organ mass) and physiological performance of hosts. Overall, our experiment is the first to show that the gut microbiome is the sole contributing factor to the adaptive physiology of a wild vertebrate host in its natural environment.

Diet is a common driver of intra- and interspecific gut microbiome variation in many taxa (humans: Turnbaugh et al., 2009; de Filippo et al., 2010; mammals: Muegge et al., 2012; Nelson et al., 2013; birds: Waite & Taylor, 2015; reptiles: Kohl et al., 2014; amphibians: Vences et al., 2016; fish: Sullam et al., 2012; insects: Jehrke et al., 2018). Variation of microbial flexibility in response to diet has also been recorded in some studies (reviewed in Hauffe & Barelli, 2019). However, most of these studies are unable to fully tease apart separate differential responses to diet from host genetics (Bolnick et al., 2014). In this study, host species as a confounding factor can be excluded since toads studied here represent populations of the same species only recently separated (< 20 years; de Villiers, 2006; Telford et al., 2019) for only approximately 6 generations (Vimercati, 2017; Vimercati et al., 2017). Therefore, our data shows that the gut microbiome can diverge its degree of microbial flexibility in response to novel conditions within a species. High microbial flexibility in invasive Cape Town toads can facilitate the rapid adjustment of microbial communities to environmental change when invasive toads spread or colonize new habitats (Wagener et al., *in review*). However, given that the other invasive populations of the same species show limited divergence of gut microbial communities from their native counterparts, compared to the invasive population investigated in this study (Wagener et al., *in review*), a high degree of microbial flexibility might, therefore, not be common in all populations of an invasive species. It is possible that the introduction of tadpoles, rather than adults, has resulted in distinct microbial communities in the Cape Town invasive toad population with a unique ability to facilitate colonisation of beneficial microbial symbionts when exposed to environmental change (Wagener et al., *in review*). However, other mechanisms, such as evolution in response to new environments, could potentially produce similar outcomes.

In many cases, hosts are dependent on their symbiotic gut microbiota to degrade complex substrates useable by the host (Bäckhed et al., 2005; Turnbaugh et al., 2006). Previous studies have found that individuals displaying divergent taxonomic responses to change are able to maintain predicted functional features (i.e. functional redundancy) (Lozupone et al., 2008; Sanders et al., 2015; Bletz et al., 2016). Contrastingly, our study found taxonomic microbial flexibility is coupled with predicted functional flexibility. Differential predicted functional features, such as increased lipid and carbohydrate metabolic pathways, in toad hosts with invasive gut microbiomes, indicate that functional flexibility allows invasive gut microbiomes to upregulate resource absorption, biosynthesis and degradation. Although we only estimated predicted functional pathways, we observed that the predicted functional flexibility was coupled with flexibility of energy investments seen in toads with invasive gut microbiomes (i.e. increased body fat % and liver mass when placed on a novel diet). Similar findings in laboratory studies with germ-free mice have demonstrated that gut microbial change is coupled with functional microbial change, but also changes in resource investments (Turnbaugh et al., 2006; Lozupone et al., 2012; Morgan et al., 2012; Heintz-Buschart & Wilmes, 2018). Microbial flexibility in wild populations can, therefore, act as a source of adaptive potential (in terms of increasing energy investment) in response to novel environmental change, as demonstrated in this study. However, future studies investigating functional metagenomic pathways will be valuable to demonstrate whether this flexibility of microbial taxa and host energy resources is coupled with the true functional potential of the gut microbiome.

Increasingly, symbiotic microbes are recognised as having a fundamental role in host phenomic plasticity (i.e. the ability of a single genotype to adjust its expression in order to display varying phenotypes) and may be influencing host adaptation to environmental change (Chevalier et al., 2015; Alberdi et al., 2016; Voolstra & Ziegler, 2020). Vertebrates’ adaptation to novel environments can be impacted by not only the interaction of hosts’ genotype with its environment but also the interaction of the hosts’ hologenome (i.e. the collection of the host and its symbiotic microbes genomes) with the hosts’ environment (Alberdi et al., 2016; Zilber-Rosenberg & Rosenberg, 2008; Shapira, 2016). A host’s gut microbial composition and/or functional gene expression can, therefore, improve the capacity of hosts to acclimate and adapt physiologically to environmental change (Alberdi et al., 2016). Within this framework, the present study’s results on host physiology (resource investment and physiological performance) provides evidence that gut microbial changes are coupled with a hosts’ ability to adapt physiologically to novel environments. A previous study has shown that invasive guttural toads outperform native individuals, possibly indicating physiological adaptation to its novel introduced environment (Vimercati et al., 2019). The results of the present study provide evidence that the gut microbiome mediates this ecological adaptation in an invasive amphibian, potentially explaining why the changes appeared so quickly (< 20 years) in this population. On the other hand, studies on plant invasions highlight enemy release (i.e., release from native pathogens in the introduced range) as a possible explanation for improved host physiology in invasive regions (Chun et al., 2010). Absence of native gut bacterial pathogens in invasive gut microbiomes could, therefore, also lead to the increased physiological performance observed in the guttural toads’ invasive region. Investigations into existing laboratory models, such as rats (*Rattus norvegicus*), the house mouse (*Mus musculus*), African clawed frog (*Xenopus laevis*) and three-spined sticklebacks (*Gasterosteus aculeatus*), hold great potential when combined with studies of their invasive populations to provide more insights into the link between host physiological adaptation and the gut microbiome.

Although our study demonstrates that gut microbiome mediates resource uptake and physiological performance, we found no effects on body mass and lean structural mass. Variation of lean structural mass and body mass between the invasive and native guttural toad populations has been reported in previous studies (Vimercati et al., 2018; Vimercati et al., 2019). It is possible that our study period (only three weeks of FMTs) was too short to induce marked changes in these organs. Long-term FMT experiments are recommended to determine whether gut microbial communities can alter other physiological attributes of the host and if FMTs can bring about long-term benefits in host physiology.

Symbiotic interactions with bacteria enhance invasions of many alien plant (Richardson & Pyšek, 2012; Traveset & Richardson, 2014) and insect species (Lu et al., 2016). However, the contributions of symbiotic bacterial communities to the success of vertebrate invasions has received little attention (see Wagener et al., *in review*). Most studies investigating host-microbial relationships in vertebrate invasions centre around the impact of pathogen loss (i.e. enemy release hypothesis) on invasive populations (Colautti et al., 2004; Phillips et al., 2010; Prior et al., 2015). It is evident, from our results, that host-associated bacterial communities can have large impacts on host adjustments or adaptation to new environments and ultimately, host fitness. Novel interactions between newly acquired symbionts and their hosts can lead to enhanced performance of invasive species and facilitate establishment in non-native areas (i.e. enhanced mutualism hypothesis; Sun & He, 2010; Coats & Rumpho, 2014). The acquisition of new symbionts in the guttural toads’ invasive region could have led to a unique invasive microbiome enhancing their hosts’ ability to respond to novel conditions both functionally and physiologically. Invasive populations are generally known to experience a lag phase between colonisation and expansion, during which time they are thought to evolve adaptations that determine invasion success (Keller & Taylor, 2008). If it takes some time to acquire novel relationships with native microbiota, invasive populations might, therefore, experience a ‘microbial lag phase’. However, it is possible that release from native pathogenic gut bacteria could have led to the observed physiological differentiation in the invasive population, as have been observed in plant invasions (Chun et al., 2010). Whether perceived host physiological variation is the result of either mutualistic, commensalistic or parasitic relationships between bacterial communities and their invasive hosts is an important point to consider when conducting future studies investigating the role of symbiotic bacteria on host responses to environmental change. Nevertheless, we highlight the imperative to identify not only the invasion potential of an introduced vertebrate population but also their microbial symbionts, as is already recognized in plants.

Both animals and plants harbour microbes that affect their physiology and subsequently fitness. Our study is the first to demonstrate that microbial symbionts are important mediators of a wild vertebrate species’ physiological responses to environmental change, with rigorous reciprocal transplant experiments. The importance of microbial communities in facilitating plant and insect invasions has been researched extensively (Traveset & Richardson, 2014; Lu et al., 2016). However, our study is the first to show that the vertebrate microbiome can impact an invasive hosts’ physiology and ultimately increase its invasion potential. In some of the most destructive invasions, the invader is not a single species but a mutualistic complex, and its invasion ecology cannot be understood without considering the interactions between the hosts and its microbial symbionts, for example *Chromolaena odorata* and *Fusarium* species spores (Mangla et al., 2008), common reed *Phragmites australis* (Nelson & Karp, 2013) and Chinese tallow *Triadica sebifera* (Yang et al., 2013). Considering the pronounced impact of gut microbial communities on host physiology and fitness demonstrated in this study and other laboratory studies, it is surprising that less than 20 studies currently consider the impact of gut microbial communities on invasive host health and physiology. Our findings emphasize the importance and unique opportunity invasive systems provide us to explore host-microbiome evolution.

## ACKNOWLEDGEMENTS

The authors would like to thank Morne du Plessis and Jan-Hendrik Keet for assistance and advice regarding bioinformatics processing of next-generation sequencing data. All authors would like to thank Christy Momberg, Erika Nortje, Sarah Davies, Suzaan Kritzinger-Klopper and Megan Mathese for logistical and administrator support. We are extremely grateful to Mike Cherry, Katy Menell and Chloë Cherry for hosting our toads in Cape Town. We thank Prof. Alex Flemming for supplying us with an ample amount of glycerol. Thanks to the NCC group for collecting toads and faecal samples in Cape Town. In particular, we would like to thank Jonathan Bell, Lulama Zibi and Nabeelah Domingo. We thank Damian van Aswegen, Carla Madelaire, Adriana Barsotti, Jaco Wagener, Johan Wagener and Mariette Wagener for assistance with field work. We thank James Baxter-Gilbert, Allan Ellis, Jan-Hendrik Keet, Anthony Herrel, Sophie von der Heyden and Julia Riley for fruitful discussions throughout the duration of this project. Thanks particularly to Morne for reading through earlier versions of this manuscript. We would also like to thank the Durban Botanical Gardens for help with collecting toads. All authors thank the Central Analytical Facilities at Stellenbosch University for the processing and sequencing of faecal samples. In particular we thank Alvera and Carel for their timely and thorough processing of the faecal material samples. Lastly, we thank the DSI-NRF Centre of Excellence for Invasion Biology (CIB) and Stellenbosch University (SU) for making our work possible.

## Notes

**Funding:** CW, NPM, and JM would like to thank the DSI-NRF Centre of Excellence for Invasion Biology (CIB) for their support.

### Competing Interest Statement

The authors have declared no competing interest.

