## Supplementary Information for "The gut microbiome facilitates ecological adaptation in an invasive vertebrate"

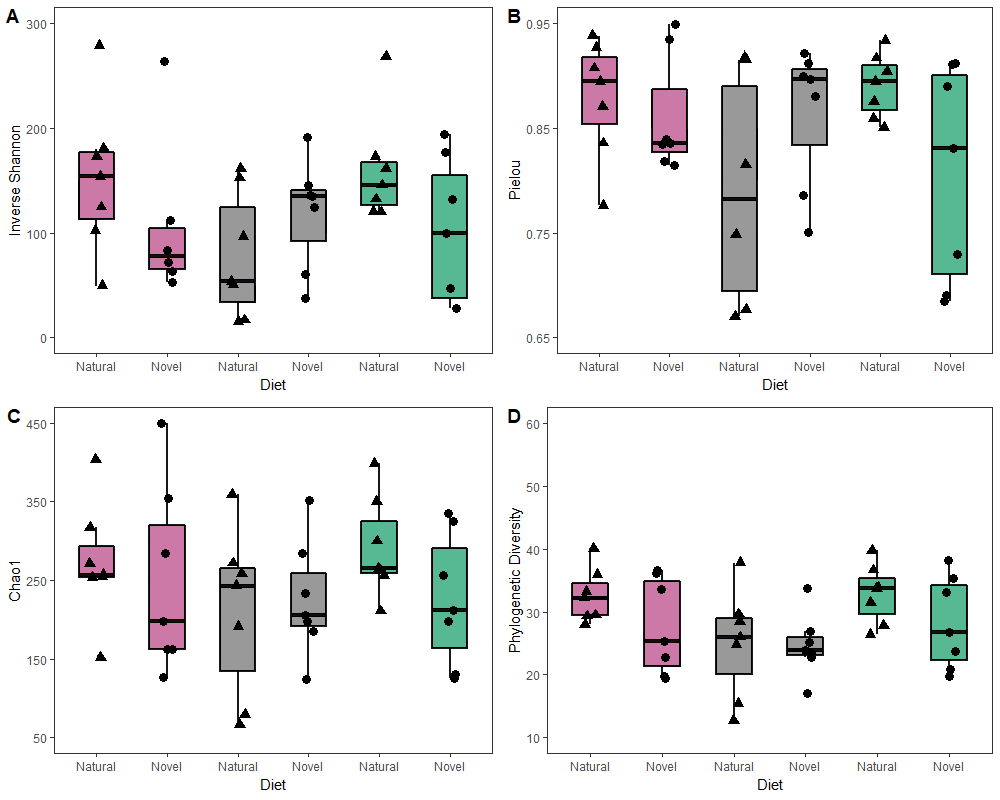
**Figure S1.** Gut bacterial alpha diversity of guttural toads (*Sclerophrys gutturalis*) on three faecal microbial treatments; native (blue), control (grey), and invasive (pink) exposed to two diets: natural or novel dietary challenge. Experiments were conducted in the native region of guttural toads, Durban, South Africa. Alpha diversity metrics (A) Shannon inverse, (B) Pielou evenness, (C) Chao1 and (D) were not significantly different between faecal microbial treatments or diets (GLMM, *p* > 0.05). The black line and whiskers in the box plots represent the medians and range of the lower quartile (25^th^ percentile) and upper quartile (75^th^ percentile).

**Figure S2.** Gut bacterial alpha diversity of guttural toads (*Sclerophrys gutturalis*) on three faecal microbial treatment groups; native (blue), control (grey), and invasive (pink) exposed to two diets; natural or novel dietary challenge. Experiments were conducted in the invasive region of guttural toads, Cape Town, South Africa. Alpha diversity metrics (A) Shannon inverse, (B) Pielou evenness, (C) Chao1 and (D) were not significantly different between faecal microbial treatments or diets (GLMM, *p* > 0.05). The black line and whiskers in the box plots represent the medians and range of the lower quartile (25^th^ percentile) and upper quartile (75^th^ percentile).

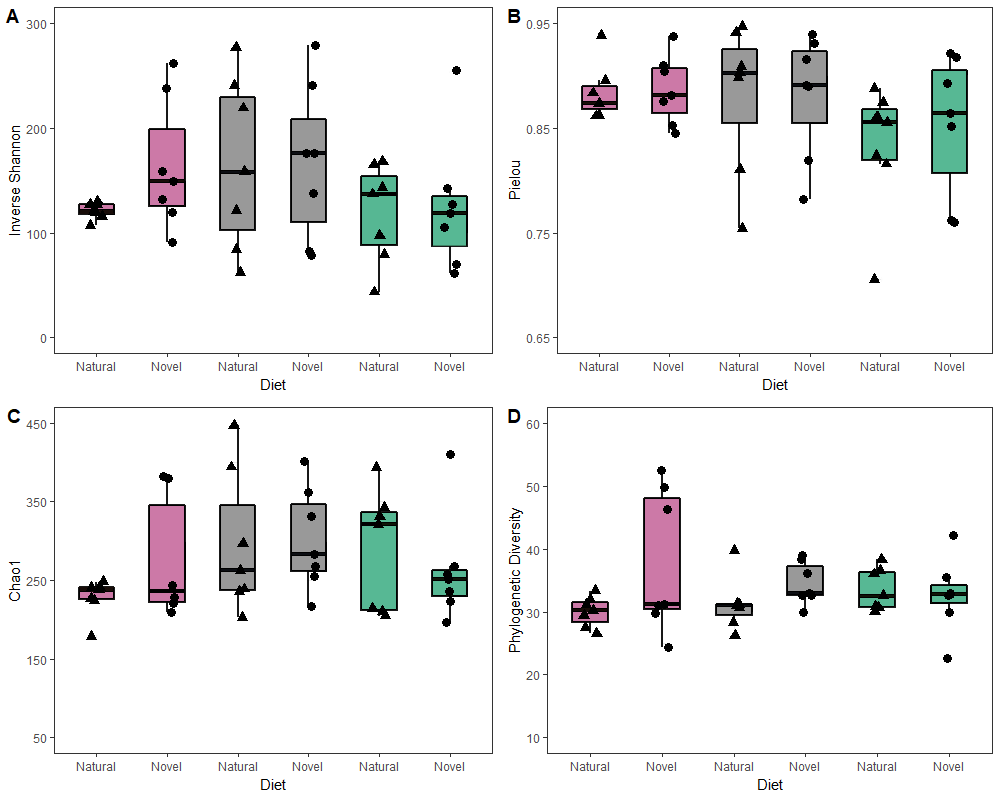

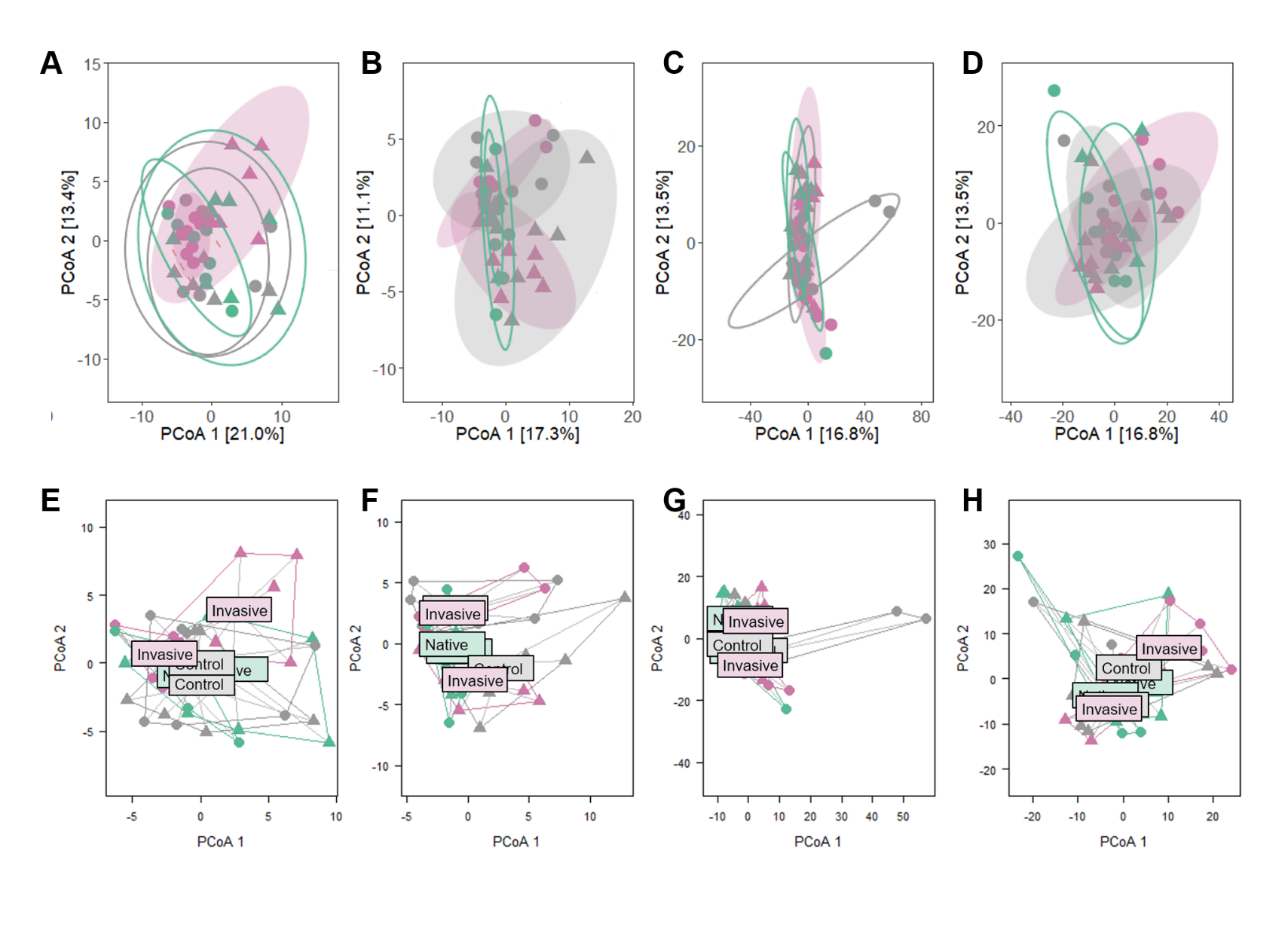
**Figure S3.** Principle Coordinates Analysis (PCoA) of PHILR-Euclidean phylogenetic (A, B, E, F) and CLR-Euclidean predicted functional (C, D, G, H) beta diversity demonstrating responses of guttural toads (*Sclerophrys gutturalis*) colonized by native (blue), invasive (pink) and control (glycerol, grey) toad gut microbial communities to natural diets (circles) and novel dietary challenge (triangles). Experiments were completed in the toads’ native range, Durban (A, C, E, G) and invasive range Cape Town, South Africa (B, D, F, H). PERMANOVA tests indicated that invasive gut microbial communities (shaded ellipses: Cape Town self-transplant, Cape Town control and Durban transplant) significantly shifted their gut microbial phylogenetic and predicted functional diversity in response to novel diets, whilst native gut microbial communities (empty ellipses: Durban self-transplant, Durban control and Cape Town transplant) showed no response to novel diets. Permutational test of dispersions (PERDISP) showed responses were not the result of variation in dispersion.

**Figure S4.** Body mass and lean structural mass of guttural toads (*Sclerophrys gutturalis*) colonized by native (blue), control (glycerol, grey) and invasive (pink) toad gut microbial communities and subsequently subjected to two diets, natural (native blue and invasive pink ants) and novel dietary challenge (yellow crickets). Experiments were completed in the toads’ native range, Durban (A, C) and invasive range, Cape Town (B, D). Body mass and lean structural mass did not vary significantly between
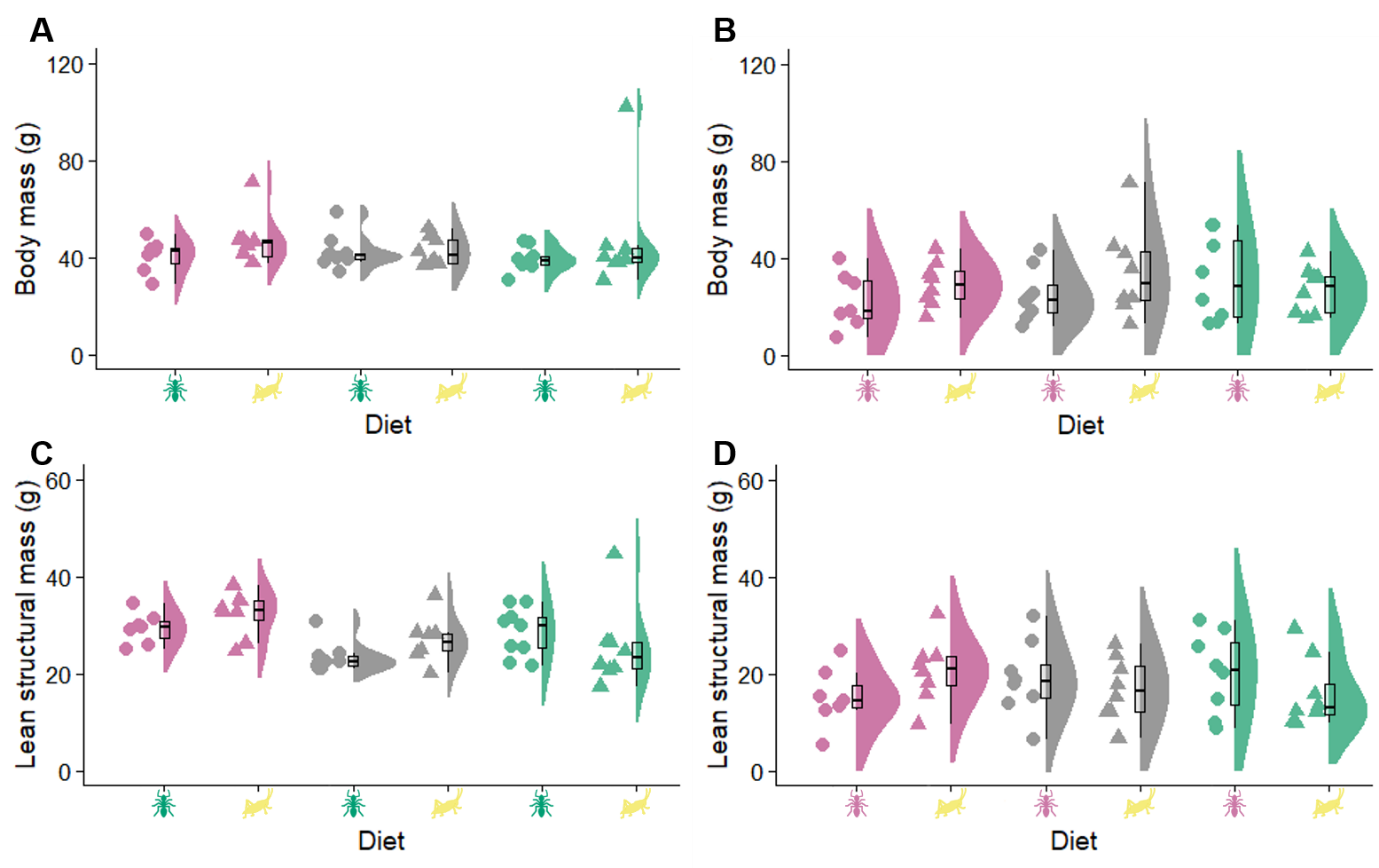
faecal microbial treatments or diets (GLM, *p* > 0.05).

**Table S1.** PERMANOVA pair-wise comparisons analysing differences between toad gut microbial composition and phylogenetic diversity as measured CLR- and PHILR-Euclidean distance matrices. Toads were subjected to either three faecal microbial treatments (FMTs); glycerol control, native faecal recipients and invasive faecal recipients and thereafter exposed to one of two diets; natural or novel dietary challenge. Experiments were conducted in the tpads’ invasive (Cape Town) and native (Durban) region. For each comparison, dependent and explanatory variables, degrees of freedom (d.f.), sum of squares (SS), pseudo-F-statistic, r-squared values(R^2^) and *p*-values are reported.

| Experimental area | Pairwise- Comparison | Dependent variable | d.f. | SS | Pseudo-F | *R^2^* | *p*-value |
| --- | --- | --- | --- | --- | --- | --- | --- |
| Cape Town | Invasive and control FMTs on dietary challenge | CLR-Euclidean | 13 | 2582.8 | 1.08 | 0.08 | > 0.05 |
|  |  | PHILR-Euclidean | 13 | 197.5 | 1.65 | 0.12 | > 0.05 |
|  | Invasive and control FMTs on natural diet | CLR-Euclidean | 13 | 2033.9 | 0.93 | 0.07 | > 0.05 |
|  |  | PHILR-Euclidean | 13 | 141.3 | 0.69 | 0.06 | > 0.05 |
|  | Invasive FMTs on natural and dietary challenge | CLR-Euclidean | 13 | 3838.2 | 1.74 | 0.13 | < 0.005 |
|  |  | PHILR-Euclidean | 13 | 399.2 | 1.91 | 0.14 | < 0.05 |
|  | Control FMTs on natural and dietary challenge | CLR-Euclidean | 13 | 3144.0 | 1.30 | 0.10 | < 0.05 |
|  |  | PHILR-Euclidean | 13 | 482.2 | 1.67 | 0.12 | < 0.05 |
|  | Native FMTs on natural and dietary challenge | CLR-Euclidean | 13 | 1471.4 | 0.94 | 0.07 | > 0.05 |
|  |  | PHILR-Euclidean | 13 | 69.8 | 0.70 | 0.05 | > 0.05 |
| Durban | Native and control FMTs on dietary challenge | CLR-Euclidean | 13 | 1188.7 | 1.00 | 0.08 | > 0.05 |
|  |  | PHILR-Euclidean | 13 | 141.4 | 1.53 | 0.11 | > 0.05 |
|  | Native and control FMTs on natural diet | CLR-Euclidean | 13 | 139.0 | 0.90 | 0.07 | > 0.05 |
|  |  | PHILR-Euclidean | 13 | 141.3 | 0.69 | 0.06 | > 0.05 |
|  | Invasive FMTs on natural and dietary challenge | CLR-Euclidean | 13 | 2175.0 | 1.79 | 0.13 | < 0.005 |
|  |  | PHILR-Euclidean | 13 | 428.9 | 2.61 | 0.18 | < 0.01 |
|  | Control FMTs on natural and dietary challenge | CLR-Euclidean | 13 | 1241.6 | 1.10 | 0.08 | > 0.05 |
|  |  | PHILR-Euclidean | 13 | 128.0 | 0.68 | 0.05 | > 0.05 |
|  | Native FMTs on natural and dietary challenge | CLR-Euclidean | 13 | 1426.4 | 1.13 | 0.09 | > 0.05 |
|  |  | PHILR-Euclidean | 13 | 261.5 | 1.69 | 0.12 | > 0.05 |

**Table S2.** Summary of best-fit mixed models analysing the gut bacterial alpha diversity differences between *Sclerophrys gutturalis* (guttural toad) on three faecal microbial treatment groups; invasive faecal recipients, native faecal recipients and controls. Toads were subsequently subjected to one of two diets; natural or novel dietary challenge. Experiments were conducted in toads’ invasive (Cape Town) and native (Durban) areas. For each model, fixed and random explanatory variables, degrees of freedom (d.f.), *chi*-square, Akaike’s information criterion (AIC), ΔAIC, marginal (*R^2^_m_*) and conditional (*R^2^_c_*) r-squared values and *p*-values are detailed.

| Experimental Area | Dependent variables | Explanatory variables | d.f. | *x*^2^ | ∆AIC | *R^2^* | *p*-value |
| --- | --- | --- | --- | --- | --- | --- | --- |
| Cape Town | Inverse Shannon | FMT treatment * diet + body condition | 35 | 4.26 | 7.19 | 0.09 | > 0.05 |
|  |  | FMT treatment + diet + body condition | 37 | 3.95 | 3.77 | 0.09 | > 0.05 |
|  |  | FMT treatment * diet | 36 | 4.37 | 5.19 | 0.10 | > 0.05 |
|  |  | FMT treatment + diet | 38 | 4.01 | 1.79 | 0.09 | > 0.05 |
|  |  | FMT treatment + body condition | 38 | 3.32 | 2.47 | 0.08 | > 0.05 |
|  |  | Diet + body condition | 39 | 0.58 | 3.37 | 0.01 | > 0.05 |
|  |  | FMT treatment | 39 | 3.42 | 0.47 | 0.08 | > 0.05 |
|  |  | Diet | 40 | 0.60 | 1.37 | 0.01 | > 0.05 |
|  |  | Body condition | 40 | 0.01 | 1.99 | 0.00 | > 0.05 |
|  |  | Null | 41 |  | 0.00 |  |  |
|  | Chao1 | FMT treatment * diet + body condition | 35 | 5.38 | 5.87 | 0.12 | > 0.05 |
|  |  | FMT treatment + diet + body condition | 37 | 2.80 | 4.92 | 0.06 | > 0.05 |
|  |  | FMT treatment * diet | 36 | 5.57 | 3.96 | 0.12 | > 0.05 |
|  |  | FMT treatment + diet | 38 | 2.70 | 3.12 | 0.06 | > 0.05 |
|  |  | FMT treatment + body condition | 38 | 1.95 | 3.84 | 0.05 | > 0.05 |
|  |  | Diet + body condition | 39 | 1.08 | 2.93 | 0.02 | > 0.05 |
|  |  | FMT treatment | 39 | 1.95 | 1.95 | 0.05 | > 0.05 |
|  |  | Diet | 40 | 0.76 | 1.21 | 0.02 | > 0.05 |
|  |  | Body condition | 40 | 0.18 | 1.82 | 0.00 | > 0.05 |
|  |  | Null | 41 |  | 0.00 |  |  |
|  | Pielou | FMT treatment * diet + body condition | 35 | 5.31 | 7.38 | 0.12 | > 0.05 |
|  |  | FMT treatment + diet + body condition | 37 | 5.22 | 3.76 | 0.11 | > 0.05 |
|  |  | FMT treatment * diet | 36 | 5.55 | 5.38 | 0.12 | > 0.05 |
|  |  | FMT treatment + diet | 38 | 5.47 | 1.76 | 0.12 | > 0.05 |
|  |  | FMT treatment + body condition | 38 | 5.10 | 1.99 | 0.11 | > 0.05 |
|  |  | Diet + body condition | 39 | 0.26 | 5.10 | 0.01 | > 0.05 |
|  |  | FMT treatment | 39 | 5.36 | 0.00 | 0.12 | > 0.05 |
|  |  | Diet | 40 | 0.20 | 3.20 | 0.00 | > 0.05 |
|  |  | Body condition | 40 | 0.12 | 3.28 | 0.00 | > 0.05 |
|  |  | Null | 41 |  | 1.41 |  |  |
|  | Phylogenetic diversity | FMT treatment * diet + body condition | 35 | 3.29 | 8.20 | 0.07 | > 0.05 |
|  |  | FMT treatment + diet + body condition | 37 | 1.36 | 6.47 | 0.03 | > 0.05 |
|  |  | FMT treatment * diet | 36 | 3.40 | 6.21 | 0.08 | > 0.05 |
|  |  | FMT treatment + diet | 38 | 1.40 | 4.48 | 0.03 | > 0.05 |
|  |  | FMT treatment + body condition | 38 | 0.30 | 5.66 | 0.01 | > 0.05 |
|  |  | Diet + body condition | 39 | 1.17 | 2.79 | 0.03 | > 0.05 |
|  |  | FMT treatment | 39 | 0.31 | 3.66 | 0.01 | > 0.05 |
|  |  | Diet | 40 | 1.14 | 0.82 | 0.03 | > 0.05 |
|  |  | Body condition | 40 | 0.01 | 1.99 | 0.00 | > 0.05 |
|  |  | Null | 41 |  | 0.00 |  |  |
| Durban | Inverse Shannon | FMT treatment * diet + body condition | 35 | 10.11 | 2.04 | 0.19 | > 0.05 |
|  |  | FMT treatment + diet + body condition | 37 | 3.61 | 4.28 | 0.08 | > 0.05 |
|  |  | FMT treatment * diet | 36 | 8.61 | 0.99 | 0.17 | > 0.05 |
|  |  | FMT treatment + diet | 38 | 3.25 | 2.55 | 0.07 | > 0.05 |
|  |  | FMT treatment + body condition | 38 | 3.31 | 3.62 | 0.07 | > 0.05 |
|  |  | Diet + body condition | 39 | 0.33 | 3.63 | 0.01 | > 0.05 |
|  |  | FMT treatment | 39 | 3.03 | 0.86 | 0.07 | > 0.05 |
|  |  | Diet | 40 | 0.28 | 1.71 | 0.01 | > 0.05 |
|  |  | Body condition | 40 | 0.09 | 1.91 | 0.00 | > 0.05 |
|  |  | Null | 41 |  | 0.00 |  |  |
|  | Chao1 | FMT treatment * diet + body condition | 35 | 5.03 | 6.41 | 0.11 | > 0.05 |
|  |  | FMT treatment + diet + body condition | 37 | 2.85 | 4.77 | 0.07 | > 0.05 |
|  |  | FMT treatment * diet | 36 | 4.88 | 4.66 | 0.11 | > 0.05 |
|  |  | FMT treatment + diet | 38 | 2.98 | 2.83 | 0.07 | > 0.05 |
|  |  | FMT treatment + body condition | 38 | 2.08 | 3.58 | 0.05 | > 0.05 |
|  |  | Diet + body condition | 39 | 0.85 | 3.07 | 0.02 | > 0.05 |
|  |  | FMT treatment | 39 | 2.28 | 1.62 | 0.05 | > 0.05 |
|  |  | Diet | 40 | 0.72 | 1.25 | 0.02 | > 0.05 |
|  |  | Body condition | 40 | 0.21 | 1.79 | 0.00 | > 0.05 |
|  |  | Null | 41 |  | 0.00 |  |  |
|  | Pielou | FMT treatment * diet + body condition | 35 | 12.95 | 1.19 | 0.23 | > 0.05 |
|  |  | FMT treatment + diet + body condition | 37 | 2.62 | 7.04 | 0.06 | > 0.05 |
|  |  | FMT treatment * diet | 36 | 11.86 | 0.00 | 0.22 | > 0.05 |
|  |  | FMT treatment + diet | 38 | 2.63 | 5.15 | 0.06 | > 0.05 |
|  |  | FMT treatment + body condition | 38 | 2.66 | 5.06 | 0.06 | > 0.05 |
|  |  | Diet + body condition | 39 | 0.15 | 5.80 | 0.00 | > 0.05 |
|  |  | FMT treatment | 39 | 2.69 | 3.16 | 0.06 | > 0.05 |
|  |  | Diet | 40 | 0.01 | 3.95 | 0.00 | > 0.05 |
|  |  | Body condition | 40 | 0.15 | 3.81 | 0.00 | > 0.05 |
|  |  | Null | 41 |  | 1.96 |  |  |
|  | Phylogenetic diversity | FMT treatment * diet + body condition | 35 | 8.79 | 4.69 | 0.20 | > 0.05 |
|  |  | FMT treatment + diet + body condition | 37 | 7.94 | 1.96 | 0.19 | > 0.05 |
|  |  | FMT treatment * diet | 36 | 10.62 | 2.70 | 0.21 | > 0.05 |
|  |  | FMT treatment + diet | 38 | 9.71 | 0.00 | 0.19 | > 0.05 |
|  |  | FMT treatment + body condition | 38 | 4.98 | 2.86 | 0.14 | > 0.05 |
|  |  | Diet + body condition | 39 | 3.76 | 3.54 | 0.09 | > 0.05 |
|  |  | FMT treatment | 39 | 6.66 | 0.93 | 0.14 | < 0.05 |
|  |  | Diet | 40 | 2.45 | 3.06 | 0.06 | > 0.05 |
|  |  | Body condition | 40 | 1.55 | 3.86 | 0.04 | > 0.05 |
|  |  | Null | 41 |  | 3.56 |  |  |

**Table S3.** PERMANOVA results analysing the effect of faecal microbial transplant FMT treatments (glycerol control, native faecal recipients and invasive faecal recipients) and dietary change on the gut microbial composition as measured by compositional CLR- and phylogenetic PHILR-Euclidean metrics in guttural toads (*Sclerophrys gutturalis*) from two experimental areas, Cape Town and Durban, South Africa. For each comparison, dependent and explanatory variables, degrees of freedom (d.f.), sum of squares (SS), pseudo-F-statistic, r-squared values (*R^2^*) and *p*-values are reported.

| Experimental Area | Dependent variable | Explanatory variable | d.f. | SS | Pseudo-F | *R^2^* | *p*-value |
| --- | --- | --- | --- | --- | --- | --- | --- |
| Cape Town | CLR-Euclidean | FMT treatment * diet | 2 | 5369 | 1.33 | 0.06 | < 0.001 |
|  |  | FMT treatment | 1 | 5129 | 1.27 | 0.06 | < 0.005 |
|  |  | Diet | 1 | 6424 | 1.69 | 0.04 | < 0.005 |
|  |  | Body condition | 2 | 2258 | 1.12 | 0.03 | > 0.05 |
|  |  | Residuals | 35 | 70806 |  | 0.81 |  |
|  |  | Total | 41 | 86986 |  | 1.00 |  |
|  | PHILR-Euclidean | FMT treatment * diet | 2 | 266 | 1.44 | 0.07 | < 0.05 |
|  |  | FMT treatment | 1 | 271 | 1.47 | 0.07 | < 0.05 |
|  |  | Diet | 1 | 185 | 2.00 | 0.05 | < 0.05 |
|  |  | Body condition | 2 | 90 | 0.97 | 0.02 | > 0.05 |
|  |  | Residuals | 35 | 3239 |  | 0.80 |  |
|  |  | Total | 41 | 4051 |  | 1.00 |  |
| Durban | CLR-Euclidean | FMT treatment * diet | 2 | 3065 | 1.56 | 0.06 | < 0.05 |
|  |  | FMT treatment | 1 | 3765 | 1.48 | 0.07 | < 0.001 |
|  |  | Diet | 1 | 1788 | 1.35 | 0.03 | < 0.005 |
|  |  | Body condition | 2 | 1626 | 1.27 | 0.06 | > 0.05 |
|  |  | Residuals | 35 | 42217 |  | 0.80 |  |
|  |  | Total | 41 | 52462 |  | 1.00 |  |
|  | PHILR-Euclidean | FMT treatment * diet | 2 | 3065 | 1.56 | 0.06 | < 0.05 |
|  |  | FMT treatment | 1 | 3765 | 1.48 | 0.07 | < 0.05 |
|  |  | Diet | 1 | 1788 | 1.35 | 0.03 | < 0.05 |
|  |  | Body condition | 2 | 1626 | 1.27 | 0.06 | > 0.05 |
|  |  | Residuals | 35 | 42217 |  | 0.80 |  |
|  |  | Total | 41 | 52462 |  | 1.00 |  |

**Table S6.** PERMANOVA results analysing the effect of faecal microbial transplant FMT treatments (glycerol control, native faecal recipients and invasive faecal recipients) and dietary change on the gut microbial predicted functional pathways as measured by compositional CLR-Euclidean metrics in guttural toads (*Sclerophrys gutturalis*) from two experimental areas, Cape Town and Durban, South Africa. For each comparison, dependent and explanatory variables, degrees of freedom (d.f.), sum of squares (SS), pseudo-F-statistic, r-squared values(R^2^) and p-values are reported.

| Experimental Area | Explanatory variables | d.f. | SS | Pseudo-F | *R^2^* | *p*-value |
| --- | --- | --- | --- | --- | --- | --- |
| Cape Town | FMT treatment * diet | 2 | 2027 | 1.57 | 0.07 | < 0.05 |
|  | FMT treatment | 1 | 1337 | 1.04 | 0.05 | > 0.05 |
|  | Diet | 1 | 1045 | 1.62 | 0.04 | < 0.05 |
|  | Body condition | 2 | 637 | 0.99 | 0.02 | > 0.05 |
|  | Residuals | 35 | 22526 |  | 0.82 |  |
|  | Total | 41 | 27572 |  | 1.00 |  |
| Durban | FMT treatment * diet | 2 | 1704 | 1.01 | 0.05 | > 0.05 |
|  | FMT treatment | 1 | 2228 | 1.38 | 0.06 | < 0.05 |
|  | Diet | 1 | 1942 | 2.41 | 0.06 | < 0.001 |
|  | Body condition | 2 | 592 | 0.74 | 0.02 | > 0.05 |
|  | Residuals | 35 | 28167 |  | 0.81 |  |
|  | Total | 41 | 34634 |  | 1.00 |  |

**Table S7.** Summary of PERMANOVA pairwise comparisons of *Sclerophrys gutturalis* (guttural toad) gut microbial predicted functional diversity as measured by CLR-Euclidean metrics. Guttural toads on three different faecal microbial treatments (invasive faecal recipients, native faecal recipients and control) was subjected to one of two diets; natural diet or novel dietary challenge. Experiments were conducted in the toads’ invasive (Cape Town) and native (Durban) region. For each comparison, dependent variable, degrees of freedom (d.f.), sum of squares (SS), pseudo-*F*-statistic, r-squared values(*R^2^*) and *p*-values are reported.

| Experimental area | Pairwise- Comparison | Dependent variable | d.f. | SS | Pseudo-F | *R^2^* | *p*-value |
| --- | --- | --- | --- | --- | --- | --- | --- |
| Cape Town | Invasive and control FMTs on dietary challenge | CLR-Euclidean | 13 | 1860.2 | 1.54 | 0.06 | > 0.05 |
|  | Invasive and control FMTs on natural diet | CLR-Euclidean | 13 | 1732.0 | 1.22 | 0.11 | > 0.05 |
|  | Invasive FMTs on natural and dietary challenge | CLR-Euclidean | 13 | 1401.0 | 2.72 | 0.18 | < 0.01 |
|  | Control FMTs on natural and dietary challenge | CLR-Euclidean | 13 | 2103.2 | 1.88 | 0.13 | < 0.05 |
|  | Native FMTs on natural and dietary challenge | CLR-Euclidean | 13 | 625.3 | 1.16 | 0.02 | > 0.05 |
| Durban | Native and control FMTs on dietary challenge | CLR-Euclidean | 13 | 1565.8 | 1.23 | 0.07 | > 0.05 |
|  | Native and control FMTs on natural diet | CLR-Euclidean | 13 | 1599.3 | 1.44 | 0.06 | > 0.05 |
|  | Invasive FMTs on natural and dietary challenge | CLR-Euclidean | 13 | 1151.9 | 1.65 | 0.11 | < 0.05 |
|  | Control FMTs on natural and dietary challenge | CLR-Euclidean | 13 | 1702.4 | 1.63 | 0.12 | > 0.05 |
|  | Native FMTs on natural and dietary challenge | CLR-Euclidean | 13 | 792.2 | 1.20 | 0.09 | > 0.05 |

**Table S10.** GLM results analysing the effect of faecal microbial transplant FMT treatments (glycerol control, native faecal recipients and invasive faecal recipients) and dietary change on the scaled body mass, scaled lean structural mass, body fat % of guttural toads (*Sclerophrys gutturalis*) from two experimental areas, Cape Town and Durban, South Africa. For each comparison, dependent and explanatory variables, degrees of freedom (d.f.), ChiSq (*x^2^*), ∆AIC, r-squared values (*R^2^_m_*) and *p*-values are reported.

| Experimental Area | Dependent variable | Explanatory variable | d.f. | *x^2^* | ∆AIC | *R^2^_m_* | *p*-value |
| --- | --- | --- | --- | --- | --- | --- | --- |
| Cape Town | Scaled body mass | FMT treatment * diet | 42 | 1.0 | 8.9 | 0.02 | > 0.05 |
|  |  | FMT treatment + diet | 44 | 0.3 | 5.7 | 0.01 | > 0.05 |
|  |  | FMT treatment | 45 | 0.1 | 3.9 | 0.00 | > 0.05 |
|  |  | Diet | 46 | 0.2 | 1.8 | 0.00 | > 0.05 |
|  |  | Null | 47 |  | 0.0 | 0.00 | > 0.05 |
|  | Scaled lean structural mass | FMT treatment * diet | 42 | 1.5 | 8.3 | 0.0 | > 0.05 |
|  |  | FMT treatment + diet | 44 | 1.2 | 4.7 | 0.0 | > 0.05 |
|  |  | FMT treatment | 45 | 0.9 | 3.0 | 0.0 | > 0.05 |
|  |  | Diet | 46 | 0.4 | 1.6 | 0.0 | > 0.05 |
|  |  | Null | 47 |  | 0.0 | 0.0 |  |
|  | Body fat % | FMT treatment * diet | 42 | 94.0 | 0.3 | 0.67 | < 0.001 |
|  |  | FMT treatment + diet | 44 | 88.0 | 0.0 | 0.65 | < 0.001 |
|  |  | FMT treatment | 45 | 9.8 | 41.2 | 0.17 | < 0.005 |
|  |  | Diet | 46 | 43.7 | 16.7 | 0.48 | < 0.001 |
|  |  | Null | 47 |  | 46.7 | 0.00 |  |
|  | Scaled liver mass | FMT treatment * diet | 41 | 52.8 | 0.0 | 0.54 | < 0.001 |
|  |  | FMT treatment + diet | 43 | 25.6 | 0.9 | 0.50 | < 0.001 |
|  |  | FMT treatment | 44 | 15.8 | 18.7 | 0.26 | < 0.001 |
|  |  | Diet | 45 | 15.7 | 17.1 | 0.25 | < 0.001 |
|  |  | Null | 46 |  | 29.1 | 0.00 |  |
| Durban | Scaled body mass | FMT treatment * diet | 42 | 1.0 | 8.9 | 0.02 | > 0.05 |
|  |  | FMT treatment + diet | 44 | 0.3 | 5.7 | 0.00 | > 0.05 |
|  |  | FMT treatment | 45 | 0.1 | 3.9 | 0.00 | > 0.05 |
|  |  | Diet | 46 | 0.2 | 1.8 | 0.00 | > 0.05 |
|  |  | Null | 47 |  | 0.0 | 0.00 |  |
|  | Scaled lean structural mass | FMT treatment * diet | 42 | 1.5 | 8.3 | 0.0 | > 0.05 |
|  |  | FMT treatment + diet | 44 | 1.2 | 4.7 | 0.0 | > 0.05 |
|  |  | FMT treatment | 45 | 0.9 | 3.0 | 0.0 | > 0.05 |
|  |  | Diet | 46 | 0.3 | 1.6 | 0.0 | > 0.05 |
|  |  | Null | 47 |  | 0.0 | 0.0 |  |
|  | Body fat % | FMT treatment * diet | 43 | 34.1 | 2.4 | 0.43 | < 0.001 |
|  |  | FMT treatment + diet | 45 | 13.1 | 0.0 | 0.42 | < 0.001 |
|  |  | FMT treatment | 46 | 18.9 | 9.4 | 0.28 | < 0.001 |
|  |  | Diet | 47 | 9.7 | 15.0 | 0.17 | < 0.001 |
|  |  | Null | 48 |  | 22.2 | 0.00 |  |
|  | Scaled liver mass | FMT treatment * diet | 41 | 52.8 | 0.0 | 0.5 | < 0.001 |
|  |  | FMT treatment + diet | 43 | 45.6 | 0.9 | 0.5 | < 0.001 |
|  |  | FMT treatment | 44 | 15.8 | 18.7 | 0.3 | < 0.001 |
|  |  | Diet | 45 | 15.7 | 17.1 | 0.3 | < 0.001 |
|  |  | Null | 46 |  | 29.1 | 0.0 |  |

**Table S11.** Summary of pairwise comparisons of *Sclerophrys gutturalis* (guttural toad) body fat % and liver mass. Guttural toads on three different faecal microbial treatments (invasive faecal recipients, native faecal recipients and control) was subjected to one of two diets; natural diet or novel dietary challenge. Experiments were conducted in the toads’ invasive (Cape Town) and native (Durban) region. For each comparison, dependent variable, degrees of freedom (d.f.), mean and standard deviation (*SD*), F-, *t*- and *p*-values are reported.

| Experimental area | Pairwise Comparison | Dependent variable | d.f. | Mean (±*SD*) | F-value | *t*-value | *p*-value |
| --- | --- | --- | --- | --- | --- | --- | --- |
| Cape Town | Invasive and control FMTs on dietary challenge | Body fat % | 15 | 0.034 (± 0.011) and 0.032 (± 0.005) | 3.90 | 0.30 | > 0.05 |
|  |  | Liver mass | 15 | 1.370 (± 0.572) and 1.530 (± 0.344) | 2.77 | -0.79 | > 0.05 |
|  | Invasive and control FMTs on natural diet | Body fat % | 15 | 0.009 (± 0.008) and 0.006 (± 0.007) | 1.11 | 1.01 | > 0.05 |
|  |  | Liver mass | 15 | 0.472 (± 0.204) and 0.753 (± 0.860) | 0.06 | -0.54 | > 0.05 |
|  | Invasive FMTs on natural and dietary challenge | Body fat % | 15 | 0.034 (± 0.011) and 0.009 (± 0.008) | 1.98 | 4.89 | < 0.001 |
|  |  | Liver mass | 15 | 1.370 (± 0.572) and 0.472 (± 0.204) | 7.88 | 4.04 | < 0.01 |
|  | Control FMTs on natural and dietary challenge | Body fat % | 15 | 0.032 (± 0.005) and 0.006 (± 0.007) | 1.98 | 6.11 | < 0.001 |
|  |  | Liver mass | 15 | 1.530 (± 0.344) and 0.753 (± 0.860) | 7.88 | 2.85 | < 0.05 |
|  | Native FMTs on natural and dietary challenge | Body fat % | 15 | 0.002 (± 0.003) and 0.011 (± 0.008) | 0.25 | 0.12 | > 0.05 |
|  |  | Liver mass | 15 | 0.451 (± 0.207) and 0.290 (± 0.218) | 0.91 | 1.66 | > 0.05 |
| Durban | Native and control FMTs on dietary challenge | Body fat % | 15 | 0.013 (± 0.009) and 0.131 (± 0.009) | 0.93 | 0.04 | > 0.05 |
|  |  | Liver mass | 15 | 1.140 (± 0.308) and 0.939 (± 0.333) | 1.17 | -1.29 | > 0.05 |
|  | Native and control FMTs on natural diet | Body fat % | 15 | 0.006 (± 0.004) and 0.007 (± 0.008) | 3.69 | 0.13 | > 0.05 |
|  |  | Liver mass | 15 | 0.859 (± 0.456) and 0.745 (± 0.126) | 0.08 | -0.41 | > 0.05 |
|  | Invasive FMTs on natural and dietary challenge | Body fat % | 15 | 0.034 (± 0.013) and 0.015 (± 0.009) | 2.06 | 3.49 | < 0.01 |
|  |  | Liver mass | 15 | 1.630 (± 0.394) and 1.110 (± 0.428) | 0.85 | 2.31 | < 0.05 |
|  | Control FMTs on natural and dietary challenge | Body fat % | 15 | 0.013 (± 0.009) and 0.007 (± 0.008) | 2.06 | 1.37 | > 0.05 |
|  |  | Liver mass | 15 | 0.939 (± 0.333) and 0.745 (± 0.126) | 6.96 | 1.43 | > 0.05 |
|  | Native FMTs on natural and dietary challenge | Body fat % | 15 | 0.013 (± 0.009) and 0.006 (± 0.004) | 4.59 | 1.53 | > 0.05 |
|  |  | Liver mass | 15 | 1.140 (± 0.308) and 0.859 (± 0.456) | 0.46 | 1.61 | > 0.05 |

**Table S12.** Summary of pairwise comparisons of *Sclerophrys gutturalis* (guttural toad) physiological performance; body fat % and liver mass (g) across three different faecal microbial transplant treatments; invasive faecal recipients, native faecal recipients and control. Experiments were conducted in the toads’ invasive (Cape Town) and native (Durban) region. For each comparison, dependent variable, degrees of freedom (d.f.), mean and standard deviation (*SD*), F-, *t*- and *p*-values are reported.

| Experimental area | Pairwise- Comparison | Dependent variable | d.f. | Mean (±*SD*) | F-value | *t*-value | *p*-value |
| --- | --- | --- | --- | --- | --- | --- | --- |
| Cape Town | Invasive and control FMTs | Body fat % | 15 | 0.02 (± 0.02) and 0.02 (± 0.02) | 0.63 | 0.12 | > 0.05 |
|  |  | Liver mass | 15 | 0.95 (± 0.63) and 1.14 (± 0.075) | 0.72 | 0.37 | > 0.05 |
|  | Invasive and native FMTs | Body fat % | 15 | 0.02 (± 0.02) and 0.01 (± 0.01) | 3.05 | 1.66 | < 0.05 |
|  |  | Liver mass | 15 | 0.95 (± 0.63) and 0.37 (± 0.22) | 4.28 | 2.65 | < 0.05 |
|  | Control and native FMTs | Body fat % | 15 | 0.95 (± 0.63) and 0.37 (± 0.22) | 3.10 | 1.75 | < 0.05 |
|  |  | Liver mass | 15 | 0.95 (± 0.63) and 0.37 (± 0.22) | 4.47 | 2.34 | < 0.05 |
| Durban | Invasive and control FMTs | Body fat % | 15 | 0.02 (± 0.01) and 0.01 (± 0.01) | 4.56 | 3.00 | < 0.05 |
|  |  | Liver mass | 15 | 1.39 (± 0.48) and 0.84 (± 0.26) | 3.48 | 2.57 | < 0.05 |
|  | Invasive and native FMTs | Body fat % | 15 | 0.02 (± 0.01) and 0.01 (± 0.01) | 4.21 | 2.51 | < 0.05 |
|  |  | Liver mass | 15 | 1.39 (± 0.48) and 0.99 (± 0.41) | 3.20 | 2.49 | < 0.05 |
|  | Native and control FMTs | Body fat % | 15 | 0.01 (± 0.01) and 0.01 (± 0.01) | 0.42 | 0.31 | > 0.05 |
|  |  | Liver mass | 15 | 0.99 (± 0.41) and 0.84 (± 0.26) | 0.99 | 0.63 | > 0.05 |

**Table S13.** GLM results analysing the effect of faecal microbial transplant FMT treatments (glycerol control, native faecal recipients and invasive faecal recipients) and dietary change on the physiological performance, total distance travelled (m) and speed (m.s^-1^) of guttural toads (*Sclerophrys gutturalis*) in Cape Town, South Africa. For each comparison, dependent and explanatory variables, degrees of freedom (d.f.), ChiSq (*x^2^*), ∆AIC, r-squared values (*R^2^_m_*) and *p*-values are reported.

|  | Explanatory variable | |  |  |  |  |  |
| --- | --- | --- | --- | --- | --- | --- | --- |
| Dependent variable | Fixed | Random | d.f. | x^2^ | ΔAIC | *R^2^_m_* | *p*-value |
| Total distance travelled | FMT treatment * diet + SVL | Trial Number | 9 | 1.8 | 2.3 | 0.48 | > 0.05 |
|  | FMT treatment * diet | Trial Number | 8 | 0.4 | 2.1 | 0.46 | > 0.05 |
|  | FMT treatment + diet + SVL | Trial Number | 7 | 1.7 | 0.4 | 0.46 | > 0.05 |
|  | FMT treatment + diet | Trial Number | 6 | 0.9 | 0.1 | 0.44 | < 0.001 |
|  | FMT treatment + SVL | Trial Number | 6 | 1.0 | 1.0 | 0.44 | > 0.05 |
|  | Diet + SVL | Trial Number | 5 | 1.0 | 26.4 | 0.02 | > 0.05 |
|  | FMT treatment | Trial Number | 5 | 26.4 | 0.0 | 0.42 | < 0.001 |
|  | Diet | Trial Number | 4 | 1.0 | 24.4 | 0.02 | > 0.05 |
|  | SVL | Trial Number | 4 | 0.0 | 25.3 | 0.00 | > 0.05 |
|  | Null | Trial Number | 3 |  | 23.4 | 0.00 |  |
| Speed | FMT treatment * diet + SVL | Trial Number | 9 | 0.6 | 1.4 | 0.60 | > 0.05 |
|  | FMT treatment * diet | Trial Number | 8 | 7.7 | 0.0 | 0.59 | < 0.001 |
|  | FMT treatment + diet + SVL | Trial Number | 7 | 0.9 | 5.7 | 0.51 | > 0.05 |
|  | FMT treatment + diet | Trial Number | 6 | 0.5 | 4.5 | 0.50 | < 0.001 |
|  | FMT treatment + SVL | Trial Number | 6 | 1.3 | 5.0 | 0.50 | > 0.05 |
|  | Diet + SVL | Trial Number | 5 | 0.5 | 32.1 | 0.07 | > 0.05 |
|  | FMT treatment | Trial Number | 5 | 27.8 | 4.3 | 0.49 | < 0.001 |
|  | Diet | Trial Number | 4 | 0.9 | 32.5 | 0.21 | < 0.001 |
|  | SVL | Trial Number | 4 | 1.9 | 30.1 | 0.06 | > 0.05 |
|  | Null | Trial Number | 3 |  | 31.3 | 0.00 |  |

**Table S14.** Summary of pairwise comparisons of *Sclerophrys gutturalis* (guttural toad) physiological performance; total distance travelled (m) and speed (m.s^-1^) across three different faecal microbial transplant treatments; invasive faecal recipients, native faecal recipients and control. Experiments were conducted in the toads’ invasive (Cape Town). For each comparison, dependent variable, degrees of freedom (d.f.), mean and standard deviation (*SD*), F-, *t*- and *p*-values are reported.

| Pairwise- Comparison | Dependent variable | d.f. | Mean (±*SD*) | F-value | *t*-value | *p*-value |
| --- | --- | --- | --- | --- | --- | --- |
| Invasive and control FMTs | Distance | 15 | 136.77 (± 6.82) and 132.31 (± 7.78) | 1.20 | 0.61 | > 0.05 |
|  | Speed | 15 | 0.06 (± 0.01) and 0.06 (± 0.01) | 1.09 | 0.87 | > 0.05 |
| Invasive and native FMTs | Distance | 15 | 136.77 (± 6.82) and 45.40 (± 2.77) | 19.89 | 4.21 | < 0.001 |
|  | Speed | 15 | 0.06 (± 0.01) and 0.04 (± 0.01) | 6.99 | 1.29 | < 0.05 |
| Control and native FMTs | Distance | 15 | 132.31 (± 7.78) and 45.40 (± 2.77) | 21.63 | 5.21 | < 0.001 |
|  | Speed | 15 | 0.06 (± 0.01) and 0.04 (± 0.01) | 8.14 | 1.11 | < 0.05 |
